# Identification of a missense ARSA mutation in metachromatic leukodystrophy and its potential pathogenic mechanism

**DOI:** 10.1101/822890

**Authors:** Liyuan Guo, Bo Jin, Yidan Zhang, Jing Wang

**Affiliations:** CAS Key Laboratory of Mental Health, Institute of Psychology, Chinese Academy of Sciences, Beijing 100101, China; Department of Neurology, Children’s Hospital of Nanjing Medical University, Nanjing, Jiangsu, 210008, China; Department of Psychology, University of Chinese Academy of Sciences, Beijing 100049, China

**Keywords:** Metachromatic leukodystrophy, ARSA gene mutation, mutation effect prediction, expression profiling, bioinformatics analysis

## Abstract

**Background:** The most common type of metachromatic leukodystrophy (MLD) is an inherited lysosomal disorder caused by recessive mutations in ARSA. The biological process of MLD disease caused by candidate pathogenic mutations in the ARSA gene remains unclear.

**Methods:** We used whole-exome sequencing (WES) and Sanger sequencing to identify the pathogenic mutation in a Chinese family. Literature review and protein three-dimensional structure prediction were performed to analyse the potential pathogenesis of the identified mutations. Overexpression cell models of wild-type and mutated ARSA genes were constructed to obtain expression profiles, and weighted gene co-expression network analysis (WGCNA), hub gene detection and protein-protein interaction (PPI) analysis were carried out to compare the biological changes caused by candidate pathogenic mutations.

**Results:** We identified an ARSA c.925G>A homozygous mutation from a Chinese late-infantile MLD patient, the first report of this mutation in Asia. According to the literature and protein structure analysis, three mutations of c.925G (c.925G>A, c.925G>T, c.925G>C) in the ARSA gene were pathogenic. The transcriptome of four ARSA overexpression cell models (c.925G, c.925G>A, c.925G>T, c.925G>C) were analysed by WGCNA, Hub genes and PPI complexes.RNA-seq and bioinformatics results indicate that the mutations at c.925G cause comprehensive molecular changes related to energy metabolism, ion binding, vesicle transport and transport.

**Conclusion:** We identified a pathogenic mutation, ARSA homozygosity c.925G>A, from a Chinese MLD family. All three mutations of c.925G in the ARSA gene are pathogenic and may cause disease by dysregulating the molecular processes of ion binding, vesicle transport and ion transport.

## Introduction

Metachromatic leukodystrophy (MLD) is an inherited lysosomal disorder caused by recessive gene mutations in ARSA and PSAP.^1–4^ It is recognized that there are 5 allelic forms of MLD (MIM ID: 250100), including late-infantile, juvenile, adult, partial cerebroside sulfate deficiency, and pseudoarylsulfatase A deficiency;^3–4^ and 2 nonallelic forms, including metachromatic leukodystrophy due to saposin B deficiency (MIM ID: 249900) and multiple sulfatase deficiency or juvenile sulfatidosis (MIM ID: 272200), a disorder that combines features of a mucopolysaccharidosis with those of metachromatic leukodystrophy. MLD is a rare disorder with an estimated birth prevalence of 1.4 to 1.8 per 100,000; there are also reports of 1/40,000 to 1/170,000.^5–6^ Although a series of potential therapies, such as haematopoietic stem cell transplantation,^7^ enzyme replacement therapy^8^ and gene therapy^9^ have been explored, currently there is no curative treatment for this disease.

The most common type of MLD disease is an autosomal recessive inherited lysosomal disorder caused by mutations in the ARSA gene, located on chromosome 22q13.33, resulting in a deficiency of the enzyme arylsulfatase A. The low activity of arylsulfatase A results in the accumulation of sulfatides in the central and peripheral nervous systems, leading to demyelination.^1^ To date more than 200 ARSA allele types have been reported as ARSA-causative variants.^10^

Whole-exome sequencing (WES) is a genomic analysis technique that captures and sequences all exons in the whole genome. It is widely used for the auxiliary diagnosis of monogenic diseases and hereditary complex diseases.^11^ Studies have recommended WES as the preferred diagnostic procedure to detect potential pathogenic genetic variants in affected individuals with genetic characteristics but without clear MLD clinical diagnosis, combined with biochemical, enzymatic or imaging studies to complete the final diagnosis.^12^

RNA sequencing (RNA-seq) is an effective method to obtain the transcriptome of life embodied in status.^13^ Besides of expression status of single genes, more complex physiological processes can be reflected via system biology approaches, such as weighted gene co-expression network analysis (WGCNA). By using WGCNA gene expression data can be transformed into co-expression modules, which provide insights into signalling networks that may be responsible for phenotypic traits of interest.^14^ Moreover, hub genes analysed from co-expression modules are quite helpful for the identification of candidate biomarkers or therapeutic targets.^15–16^ Combined with protein-protein interaction (PPI) information, co-expression modules also provide valuable clues to reveal the dynamic protein complexes. Protein complexes are of great importance for understanding the principles of cellular organization and function.^17^

In current study, we reported a Chinese family with a 2-year-old boy who could not sit or walk independently. By using WES technology we identified a missense mutation in exon 5 of the ARSA gene (c.925G>A homozygous mutation) in the proband and supported the clinical diagnosis of MLD. The mutation site was first reported in Asia, a systematics literature review and bioinformatics analyses were carried out to estimate its pathogenicity. A series of ARSA-mutant cell models were consequently constructed to simulate the cell state of homozygous-mutant patients and RNA-seq and system biology analysis were performed to explore the potential molecular pathogenesis of the identified mutation.

## Methods

### Patients

The current study was approved by the Ethics Committee (NO. 201801001-1) of the Children’s Hospital Affiliated with Nanjing Medical University (Nanjing, China). Prior to the study, written informed consent for genetic tests and publication of the case details was obtained from all adult participants and the parents or the legal guardians of children involved in the study.

### Mutation analysis

Peripheral blood samples were obtained from patients and members of their families with informed consent. DNA was prepared according to standard protocols (Cat. No. CW2087, CoWin Bioscience, China).

WES was performed on the genomic DNA of the proband and members of his family (exome sequencing and bioinformatics analysis of sequencing raw data completed in Oumeng V Medical Laboratory Co., Ltd.). One microgram of genomic DNA from peripheral blood mononuclear cells (PBMCs) was used to establish the exome library. Genomic DNA was randomly sheared by sonication and then hybridized to the Agilent SureSelect Human All Exon V6 Library to enrich exonic DNA in each library, following the manufacturer’s instructions. An enriched library targeting the exome was sequenced on the Illumina HiSeq 2000 platform to generate paired-end 150 bp sequences (PE150). The mean coverage of each exon was more than 100×, allowing for the examination of the selected region and sufficient depth to accurately match 99% of the targeted exome. Variant calling was performed with reference to UCSC hg19reference genome (http://genome.ucsc.edu/).^18^ Single-nucleotide polymorphisms (SNPs) and small insertions or deletions (indels) were sifted using ANNOVAR (2015Dec14) with the defined parameters after the deletion of duplicated reads.^19^ Short insertions or deletions (indels) altering coding sequences or splicing sites were also identified by GATK (3.3.0). The common variants and non-pathogenic candidate variants were filtered against several public databases, including dbSNP (http://www.ncbi.nlm.nih.gov/projects/SNP/snp_summary.cgi) and the 1000 Genomes Project (http://www.1000genomes.org/). Multiple alignment of the protein sequence was performed across different species by the Basic Local Alignment Search Tool (http://blast.stva.ncbi.nlm.nih.gov/Blast.cgi). Online tools including Polymorphism Phenotyping version 2 (PolyPhen-2, http://genetics.bwh.harvard.edu/pph2/),^20^ Sorting Intolerant from Tolerant (SIFT, http://sift.jcvi.org/; score less than 0.05 indicates it is deleterious)^21^ and MutationTaster (http://www.mutationtaster.org/)^22^ were used to evaluate whether amino acid substitutions affected protein function.

Variants were identified as nonpathogenic and removed if they fit any of the following criteria: (1) variants with an allele frequency greater than 5%; (2) variants located in introns and not affecting splicing; (3) synonymous variants without affecting splicing; or (4) variants located in the 5’ or 3’ untranslated region. We reviewed the functions and associated OMIM phenotypes of those genes (https://omim.org/).

Sanger sequencing was used to validate candidate variants (completed in Oumeng V Medical Laboratory Co., Ltd.). Locus-specific primers used for PCR amplification and direct sequencing were designed using the online Primer3 program (http://primer3.ut.ee/). Then Sanger sequencing of PCR products was performed to validate the potential disease-causing variants with an ABI3500 sequencer (Applied Biosystems Inc., Foster City, CA, USA).^23^

### Literature review and effect prediction of ARSA c.925 mutations

A systematic literature review was performed to obtain published ARSA c.925 mutations in MLD. The detail process of literature collection and data extraction were shown in figure S1. Amino acid conservation of ARSA protein in different species was performed by using MEGA5 and DNAMAN software. All amino acid sequences used in the conservation analysis were obtained from NCBI Web site (http://www.ncbi.nlm.nih.gov/). The effect of the amino acid changes in ARSA at p.E309 was predicted by using the web server SWISS-MODEL (https://www.swissmodel.expasy.org).

### ARSA expression/mutation vector construction

The human wild-type ARSA cDNA (cloned in the pCMV6 plasmid) was purchased from Origene (Cat. No. RC204319, Origene, USA). The c.925G mutations (c.925G>A, c.925G>T, and c.925G>C, as shown in figure S2) were introduced into the ARSA plasmid by using Site-Directed Mutagenesis Kit (Cat. No. E0552s, NEB, USA). The sequences of mutated cDNA vectors were confirmed using the ABI3500 sequencer (Applied Biosystems Inc., Foster City, CA, USA).

### Cell culture and transfection

HEK293 cells (CRL-1573, ATCC, VA, USA) were cultured in high-glucose DMEM (Cat. No. 11965092, Life Technologies, USA) supplemented with 10% FBS at 37 and 5% CO_2_. Cells were seed in 12-well at an initial density of 2 x 10^5^ cells per well 24 hours before transfection and were collected 24 hours after the transfection. The transient transfection were performed by using FuGene HD transfection reagent (Cat. No. E2311, Promega, USA).

### Total RNA extraction and RNA sequencing

Total RNA was extracted from HEK293 cells via the RNApure Tissue/Cell Kit (Cat. No. CW0560, CoWin Bioscience, China). RT-qPCR was performed to validate the overexpression of ARSA gene by using SYBR Green Master Mix (Cat. No. CW2999, CoWin Bioscience, China). The expression levels of ARSA genes (forward primer: 5’-CTGCTTGAAGAGACGCTGGT-3’, reverse primer: 5’-AAGGTGACATTGGGCAGTGG-3’) were calculated using the ΔΔCt method and normalized to the Ct values of the GAPDH gene (forward primer: 5’-GACAGTCAGCCGCATCTTCT-3’, reverse primer: 5’-TTAAAAGCAGCCCTGGTGAC-3’).

RNA sequencing and subsequent bioinformatics analysis were completed by the Oumeng V Medical Laboratory. RNA sequencing was performed using the same protocol with poly-A selection of mRNA following the manufacturer’s instructions (Illumina TruSeq). Paired-end 150 bp sequencing, which was performed on Illumina HiSeq 2000 instruments, generated more than 6G clean bases data for each sample. The raw reads were quality checked using FastQC. Low-quality bases were trimmed with Trimmomatic (parameters TRAILING: 3 and SLIDINGWINDOW: 4:15).^24^ HISAT2 was used to compare clean reads to the reference genome (GRCh37).^25–26^ Then sequence mapping data (SAM) were converted into bam files and sorted according to mapped chromosomal location using SAMtools.^27^ The software StringTie was used to calculate the gene expression and outputted the standardized TPM format expression data of all genes in each sample.^28^

### Construction of WGCNAs and detection of hub genes

Expression profiling of all samples were utilized as input data to construct co-expression networks using the weighted gene WGCNA R package.^29^ The pickSoftThreshold function was used to obtain a proper soft-thresholding power (β=3), where the corresponding Scale-free Topology Fit Index was above 0.9. We calculated the Pearson’s correlation between modules and cell treatments to identify the relevant module. In general, the module whose absolute correlation ranked first among all the selected modules was considered the one related to phenotype. In each co-expression module, genes were ranked according to four ranking methods in Centiscape2.2 (Degree, stress, Closeness, and Radiality), the overlaps of the first 30 genes of each ranking method were defined as the hub genes. The modules were defined as key modules when they had a correlation >0 and p-value<0.05 with at least one of the four types of cell (c.925G, c.925G>A, c.925G>T, c.925G>C) and used for subsequent analysis.

### Function annotation and enrichment analysis

To identify biological processes and pathways that are significantly enriched by the genes in selected modules different phenotypes, the gene list of the key modules was analyzed using the Gene Functional Classification Tool (DAVID) v6.8, which is an online tool for functional annotation and enrichment analysis to reveal biological features related to large gene lists.^30^ The online tool Metascape was used to perform annotations of the hub genes from the Gene Ontology (GO) database and Kyoto Encyclopedia of Genes and Genomes (KEGG) database.^31^

### Identification of protein-protein interaction networks

The gene lists from selected modules to build a protein-protein interaction (PPI) network based on BioGrid6, InWeb_IM7 and OmniPath database in online tool Metascape. Input correct species (H. sapiens), and choose “Express Analysis” for analysis. Moreover, if the network contains between 3 and 500 proteins, the “Molecular Complex Detection” (MCODE) algorithm has been applied to identify densely connected network components. All network diagrams are generated by Cytoscape.^32^

## Results

### Clinical finding

The proband, a 27-month-old boy, was unable to stand or walk independently, had decreased lower limb muscle strength and muscle tension, and had impaired language function. He was the second boy of a healthy, nonconsanguineous couple. Before he went to the Neurology Department of Nanjing Children’s Hospital, he received physical therapy for walking disability.

The proband showed decreased lower limb muscle strength and muscle tension and electrophysiological changes of multiple peripheral neurogenic lesions on EMG. Abnormal white matter symmetry signals in bilateral cerebral hemispheres were observed in MRI images of the proband (figure 1A).

**Figure 1.**
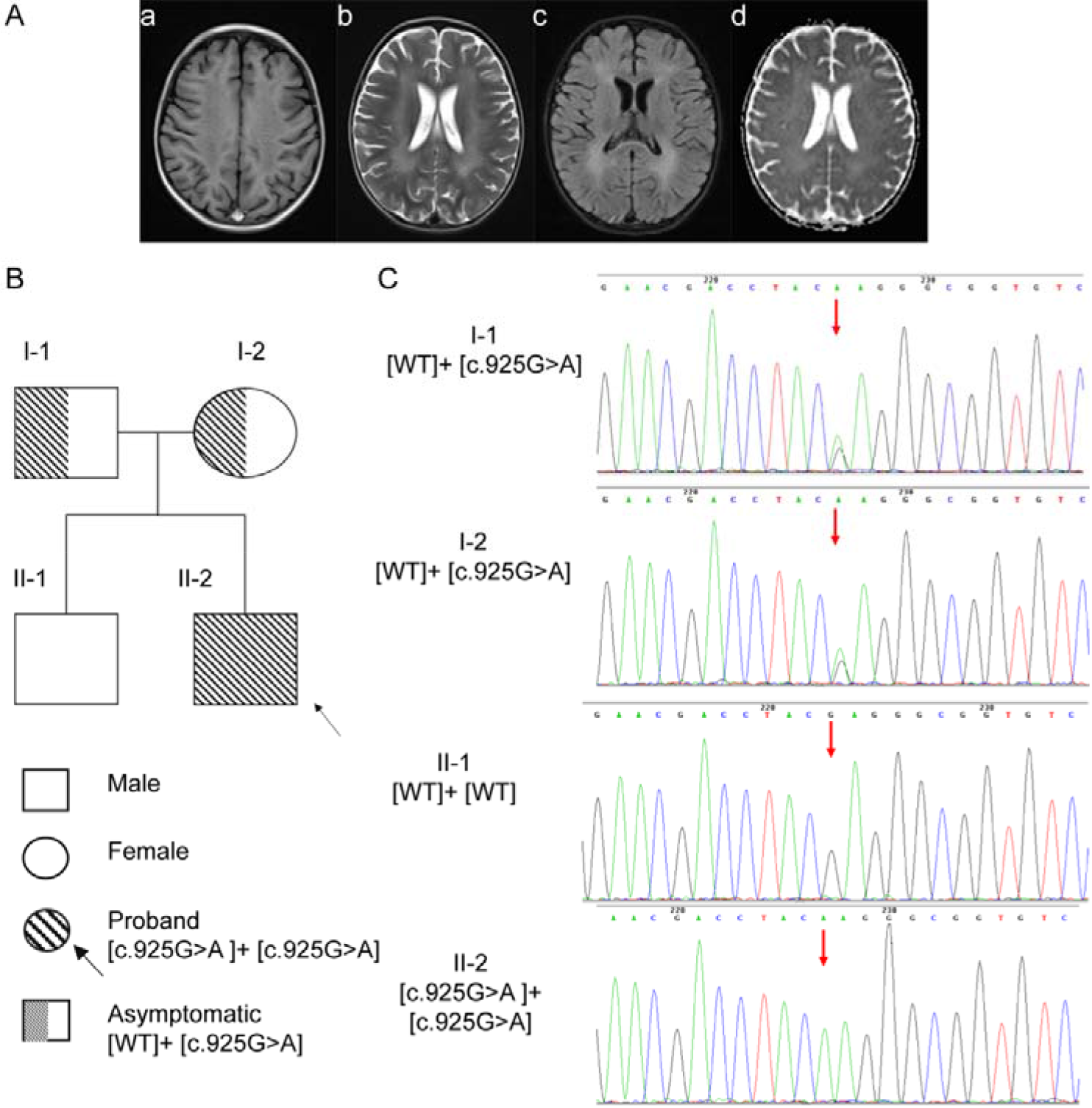
MRI from the proband and mutational analysis of the arylsulfatase A (ARSA) gene in the family. (A) MRI from the proband (II-2). Magnetic resonance imaging (MRI) shows symmetrical deep lesions located in periventricular white matter, which was low signal in T1WI (a), high signal in T2WI (b), low signal in in T2WI (c) and ep2d (d) from the proband (II-2). (B) Pedigree of the family with MLD patients. The proband was shown in the second generation with the numbers II-2. The parents of proband are first generation with the number I-1 and I-2. The healthy older brother of proband is in the second generation with the number II-1. (C) Mutational analysis of the arylsulfatase A (ARSA) gene. Genotypes of the proband shows a homozygous c.925 G>A and a heterozygous c.925 G>A in his parents. Healthy brothers do not inherit this mutation. Nucleotide numbers are derived from cDNA ARSA sequences, GenBank accession numbers: NM_000487.5 and NP_000478.3.

### Genetic Finding

We performed exome sequencing of the proband (II-2, figure 1B) and other family members in a Chinese Han family. We generated more than 17 billion Raw Bases for all samples (I-1 17.64 billion bp, I-2 19.46 billion bp, II-1 21.53 billion bp, II-2 24.75 billion bp, supplementary table S1). Among the Raw Bases, clean reads rate more than 92% (I-1 92.48%, I-2 94.9%, II-1 21.53 94.93%, II-2 94.64%, supplementary table S1) and capture efficiency more than 50% (I-1 62.25%, I-2 57.18%, II-1 63.78%, II-2 63.1%, supplementary table S1). Mapped to the targeted bases with a mean coverage of more than 100-fold (I-1 102.74-fold, I-2 103.72-fold, II-1 124.75-fold, II-2 138.26-fold, supplementary table S1). In the proband, 51535 genetic variants, including 8785 nonsynonymous variants, were identified in either coding regions or splice sites. The results of the WES test and quality control for all family members are shown in supplementary table S1-S3. According to the American College of Medical Genetics and Genomics (ACMG) recommendations, which provides interpretative categories of sequence variants and an algorithm for interpretation,^33^ it a missense mutation, c.925G>A in exon 5 of the ARSA gene (figure 1C, supplementary table S4) was identified as a candidate pathogenic locus. Genotypes of the proband showed a homozygous c.925G>A mutation, and those of the parents showed a heterozygous c.925G>A mutation (I-1, I-2, figure 1C). His healthy brother did not inherit this mutation (II-1, figure 1C).

### ARSA mutation (c.925G>A, c.925G>T, c.925G>C) function

c.925G>A (p.E309K) is a highly conserved in multiple species sequences, suggesting its structural and functional importance (figure 2A). This mutation was predicted to affect the protein features and be disease-causing by SIFT, PolyPhen2 and MutationTaster (supplementary table S4). The 3D structural model analysis showed that glutamate at position 307 was the part in the formation of a disulfide bond. Hence, three missense mutations (c.925G>A, c.925G>T, c.925G>C) caused amino acid changes (p.E309K, p.E309* and p.E309Q) and truncated the protein, altering the local charge to prevent the correct positioning of a sulfate group of the substrate and consequently its hydrolysis (figure 2). The mutation p.E309* resulted in early termination of translation (figure 2C).

**Figure 2.**
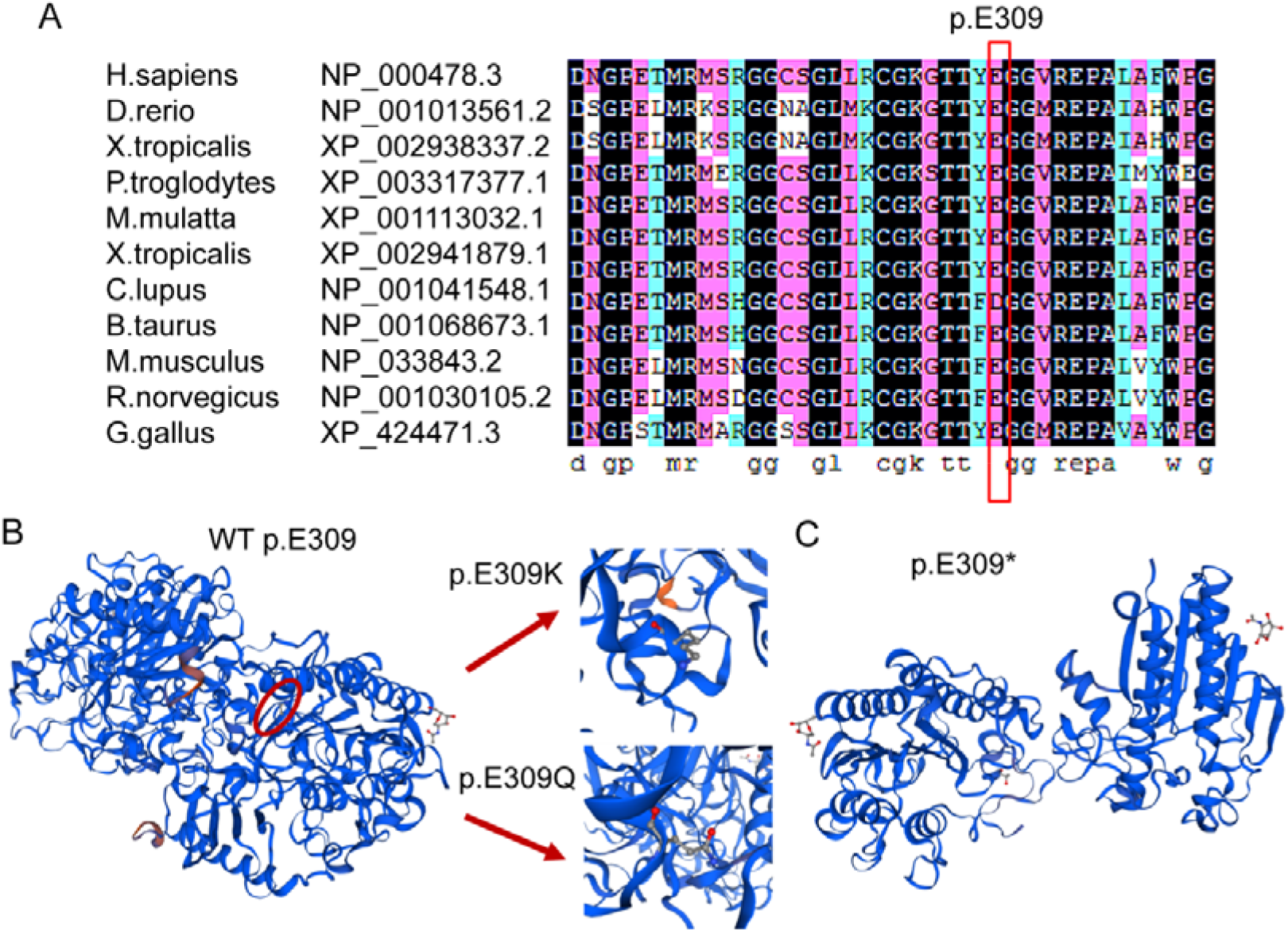
Multiple sequence alignment (MSA) and 3D structure of ARSA. (A) Multiple sequence alignment showing the sequence alignment of a specific amino acid, and its conservation in other ARSA orthologs (across different species). Nucleotide numbers are derived from cDNA ARSA sequences, GenBank accession numbers: NM_000487.5 and NP_000478.3. (B) Conformational changes induced by the p.E309K and p.E309Q missense mutation in the ARSA protein. (C) Conformational changes induced by the p.E309*, this mutation results in early termination of codons and truncated proteins.

In previous studies, ARSA c.925G (c.925G>A, c.925G>T, c.925G>C) has been reported to cause late-infancy MLD. As shown in Table 1, one of the 10 reported patients with MLD disease had adult MLD, and one was diagnosed with juvenile MLD; two patients have had c.925G>A homozygous mutations (including the proband found in this study), and other patients had composite heterozygous mutations at this site.^10 34–39^

**Table 1.** Reported summary of characteristics about the ARSA c.925G mutation Patients

| Pt No. | Fam | MLD variant | Nat/ Ethnicity | Gender | Age |  | Symptoms at onset | Genotype | Enzymatic activity | Patients' references |
| --- | --- | --- | --- | --- | --- | --- | --- | --- | --- | --- |
|  |  |  |  |  | Onset | Death |  |  |  |  |
| 1 | - | LI | Caucasian | F | 1y6m | - | Walking difficulties, dysarthria, and spasticity | [c.925 G>A ] + [c.925 G>A] | 6 % | Martina et al. (2015) |
| 2 | - | A | Caucasian | M | 40y | - | Peripheral neuropathy | [c.869 G>A] + [c.925 G>C] | - | Martina et al. (2015) |
| 3 | - | LI | Asian/Chinese | F | 4y | - | Psychomotor deterioration, motor regression, Dysarthria, Symmetrical deep WM abnormalities | [c.925 G>A] + [c.427 T>C] | - | Li et al. (2018) |
| 4 | - | LI | Asian/Chinese | F | 0y07m | - | Motor retardation, Peripheral neuropathy, Symmetrical deep WM abnormalities | [c.925 G>A ] + [c.302 G>T ] | - | Li et al. (2018) |
| 6† | Sibling | LI | European | M | 01y06m | - | Walking difficulties, , Pain attacks, Peripheral neuropathy | [c.304delC] + [c.925 G>A] | 12% | Kreysing et al. (1993) |
| 7† | - | normal | European | F | - | - | Healthy | [WT] + [c.925 G>A] | - | Kreysing et al. (1993) |
| 8† | Sibling | normal | European | M | - | - | Healthy | [WT] + [c.925 G>A] | - | Kreysing et al. (1993) |
| 9 | - | J | European | - | - | - | Progressive and profound motor deficit | [c.418 C>G] + [c.925 G>A] | 0% | Biffi et al. (2008) |
| 10‡ | Sibling | LI | Asian/Chinese | M | 01y05m | - | Walking difficulties, Peripheral neuropathy | [c.925 G>A]+ [c.925 G>A] | - | This study |
| 11‡ | - | normal | Asian/Chinese | M | - | - | Healthy | [WT] + [c.925 G>A] | - | This study |
| 12‡ | - | normal | Asian/Chinese | F | - | - | Healthy | [WT] + [c.925 G>A] | - | This study |
| 13 | - | LI | Italian | - | 02y00m | - | Spastic paraparesis Ataxia,<br>Mental deterioration,<br>Symmetrical deep WM<br>abnormalities | [c.925 G>T]+[c.59 C>A] | - | Grossi et al. (2008) |
| 14 | - | LI | Turkish | M | 01y06m | 03y06m | Difficulty in walking,<br>intentional tremor,<br>nystagmus, spontaneous<br>contraction at extremities<br>and wheezing. | [c.925 G>A]+[ c.1178 C>G] | - | Onder et al. (2009) |
| 15 | - | LI | Turkish | F | 02y06m | 04y00m | Difficulty in walking,<br>positive bilateral Babinski,<br>tremor in hands and<br>mild spasticity | [c.925 G>A]+[ c.1178 C>G] | - | Onder et al. (2009) |

Pt, patient; LI, late infantile; J, juvenile; A, adult; Presymp, presymptomatic; Fam, familiarity; F = Female; M = Male; m = month; y = year.
Sibs, sibling;
The symbol “-” indicates data not available.
Nucleotide numbers are derived from cDNA ASNA sequences, GenBank accession numbers: NM\_000487.5 and NP\_000478.3.
Mutations are described according to current mutation nomenclature guidelines (http://www.hgvs.org/mutnomen; den Dunnen and Antonarakis, 2001), ascribing the A of the first ATG translational initiation codon as nucleotide +1.
† DNA was isolated from fibroblasts of the patient No. 6, his mother No.7, and his brother No.8.
‡ DNA was isolated from peripheral blood of the patient No. 10, his father No.11 and his mother No.12.

### Transcriptome and biological analysis

Mock vector (Vector), wild-type ARSA cDNA plasmid (c.925G), and missense mutated ARSA cDNA plasmid (c.925G>A, c.925G>T, c.925G>C) were transfected into HEK293 cells, RNA-seq and consequent bioinformatics analyses were performed on untreated and transfected cells (figure 3A). As validated by RT-qPCR, the expression level of ARSA exceeded 20,000-fold in wild-type ARSA cDNA plasmid-transfected c.925G and three missense mutation-transfected (c.925G>A, c.925G>T, c.925G>C) cell models over WT and Vector cells (figure 3B).

**Figure 3.**
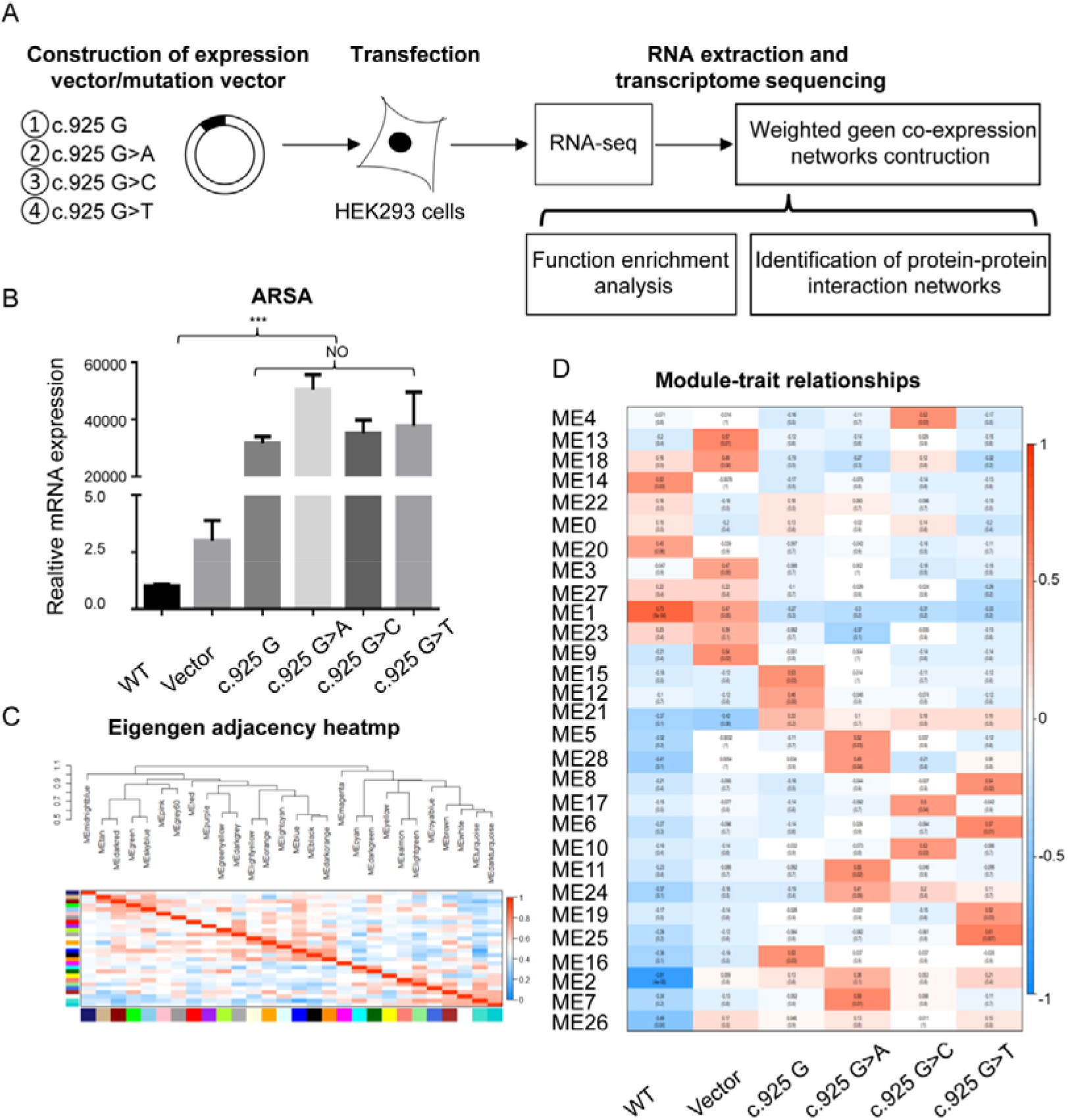
ARSA-Express/Mutation cell mode construction and biological function analysis. (A) Construction of ARSA Gene Express/Mutation cell model. (B) Relative mRNA expression of ARSA gene, GAPDH was used as a loading control. \**p*< 0.05 in independent Student *t* test, \*\*\**p*< 0.001 in independent Student *t* test). (C) Hot Map of Characteristic Gene Adjacency in Characteristic Gene Network. Each row and column corresponds to a characteristic gene (marked with the same color). In heatmap, red denotes high adjacency (positive correlation) and blue denotes low adjacency (negative correlation), as shown in the color legend. (D) Characteristic gene module-trait Association of each module. Each row corresponds to a module characteristic gene, and each column corresponds to a cell type (trait). Each cell contains Pearson correlation coefficients (numbers outside parentheses) and associated *p* values (numbers inside parentheses). According to the color legend, color coding is carried out by correlation. Red indicates positive correlation and blue indicates negative correlation.

An average of 25,280,291 raw reads (a maximum of 21,591,207 and a minimum of 32,554,987) were generated for each samples by RNA-seq. After removing low-quality and adapter sequences, an average of 25 million clean reads were obtained in each sample. The average GC content percentages in the samples ranged from 50.67% to 52.42%. The Q30 percentages were more than 93.14% (supplementary table S5). After the low-expression genes (TPM=0) had been filtered out from all gene expression libraries, 43450 genes were used for WGCNA analysis.

Twenty-nine co-expression modules were identified in the WGCNA analysis, including the grey module (ME0). According to the description of WGCNA package, genes in grey were not included in any module, so the subsequent analysis did not include them. The interactions among these 28 co-expressed modules (reflected by the connectivity of eigengenes) was shown as a heatmap of module eigengene adjacency (figure 3C). Then the module-trait relationships with all traits were calculated by correlating the module’s eigengenes to the traits of interest (figure 3D) and were used for selection of modules for downstream analysis. The module-trait relationships showed that several modules were positively correlated with one or more cell types, and it also showed a clear distinction between the different cell types. After eliminating pseudogenes and lncRNA, 14, 16, 18 and 14 hub genes were identified for c.925G, c.925G>A, c.925G>T and c.925G>C, respectively, by overlap of the top 30 genes according to four ranked methods in Centiscape2.2. These hub genes are annotated to be involved in processes such as Golgi vesicle-mediated transport, cell junction, calcium binding and carbohydrate metabolism (supplementary table S6).

Functional enrichment analysis was conducted on the gene lists of 14 modules that were significantly positively correlated with four groups of samples (c.925G, c.925G>A, c.925G>T, c.925G>C) using the DAVID Gene Functional Classification Tool. The recommended threshold enrichment score (>1.3) was to select enriched clusters, and 16 clusters were found in four groups (Table 2).

**Table 2.**
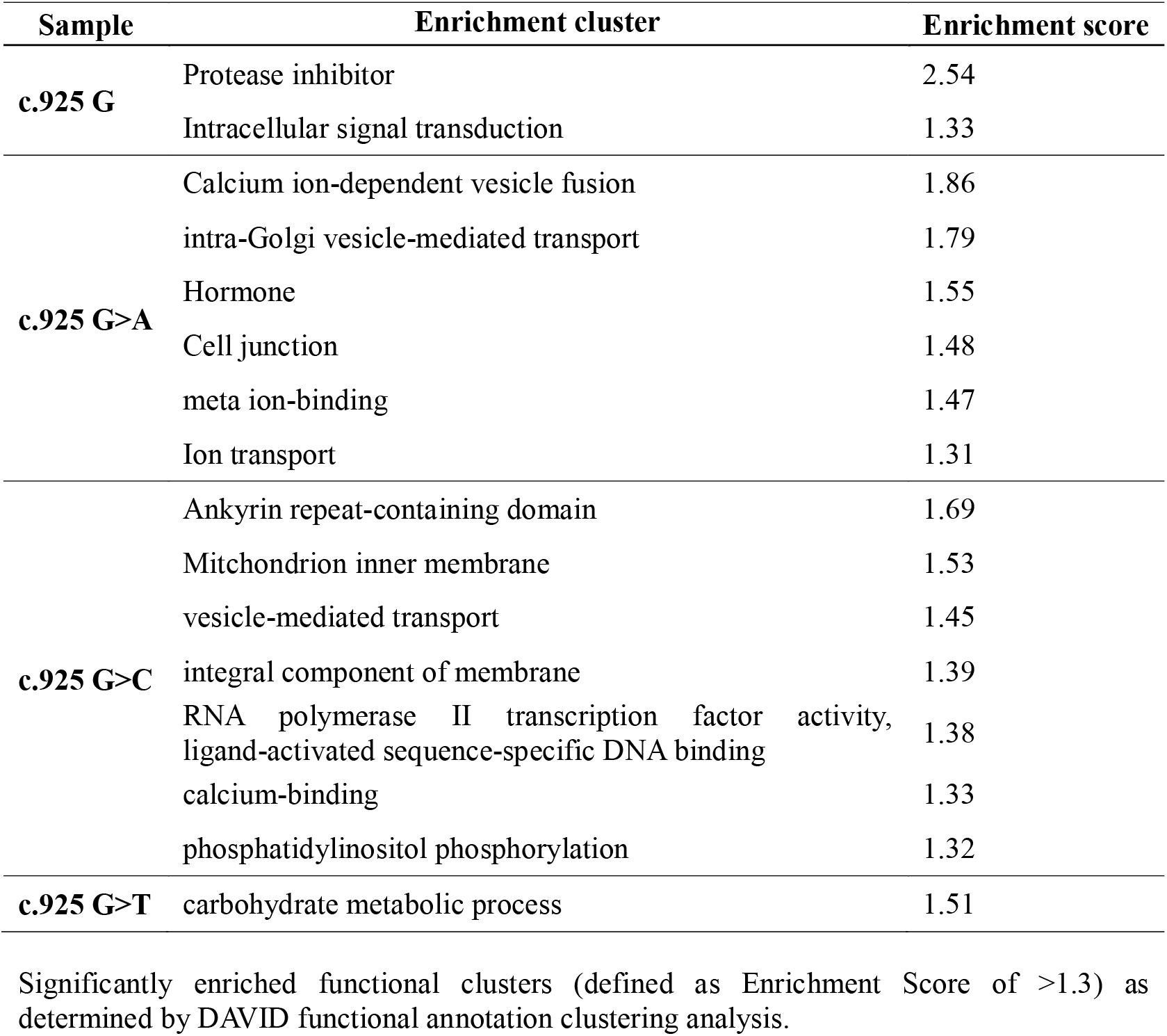
Functional enrichment analysis

The Molecular Complex Detection (MCODE) algorithm of Metascape software was applied to identify densely connected network components. 33 MCODE components were identified in the 14 modules of the four phenotypes (c.925G, c.925G>A, c.925G>T, c.925G>C). In the network of protein interactions that was identified, the mutation of c.925G (c.925G>A, 11 components; c.925G>T, 6 components; c.925G>C, 14 components) had more protein complexes that might interact than the control group (c.925G, 3 components) (figure 4).

**Figure 4.**
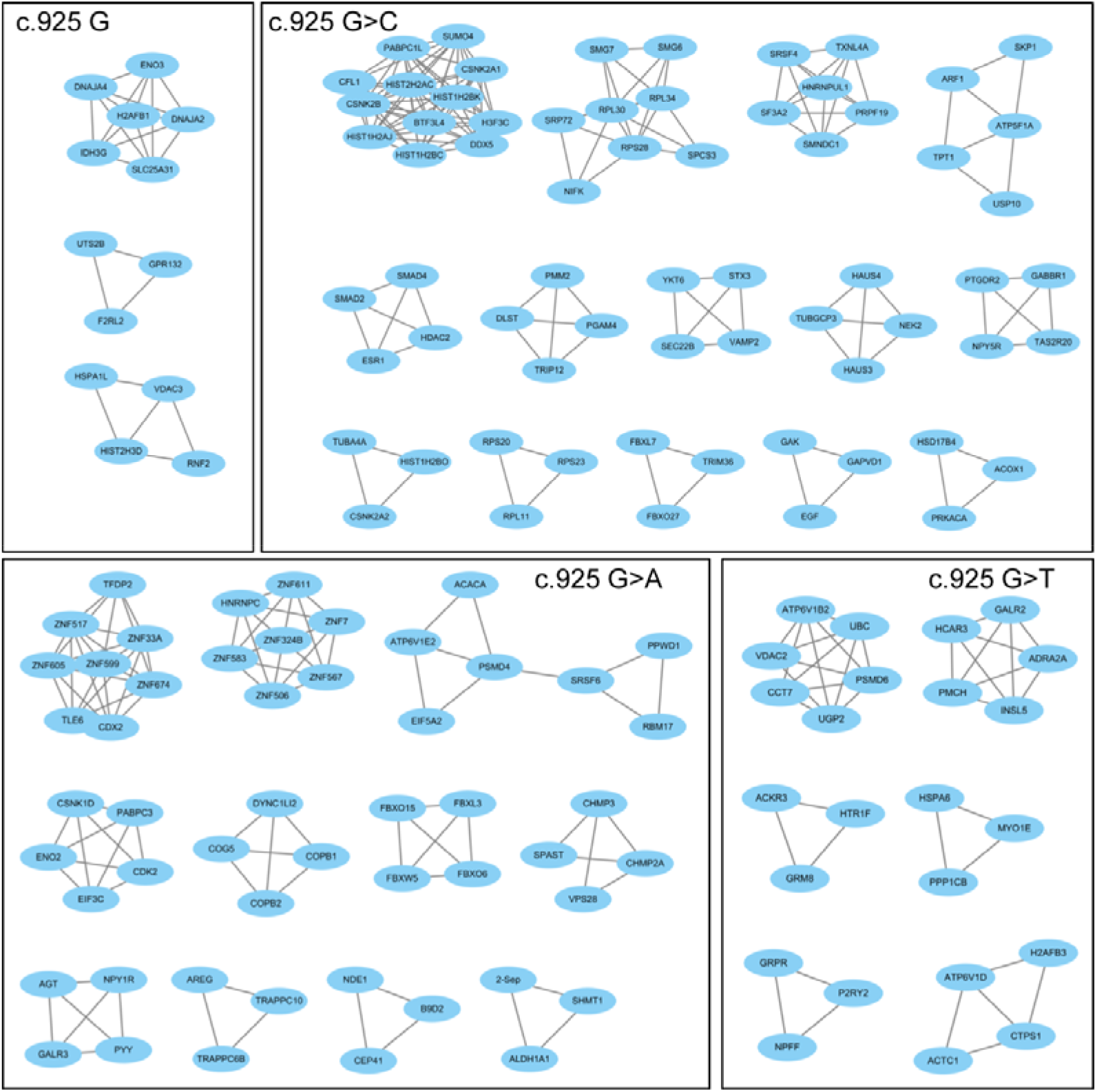
Identification of protein-protein interaction networks. The MCODE components identified in the gene list of the relevant modules corresponding to each sample.

## Discussion

We report a boy initially manifested as inability to stand alone, decreased lower limb muscle strength and muscle tone, and electrophysiological changes in multiple peripheral neurogenic lesions on the EMG. The 2-year-old male child could not sit for a long time, and his language function was impaired. The MRI results showed white matter lesions. The results of the WES test showed that an ARSA c.925G>A homozygous mutation site carried by the child was inherited from his parents. The child diagnosed with MLD disease, from an unaffected family. The ARSA gene c.925G>A mutation was first reported by the Kreysing research team. The team found that the c.304del C and c.925G>A mutation site was pathogenic, while heterozygous mutation site c.925G>A had no typical clinical manifestation in the parents.^40^ It has been found that c.925G>A is a complex heterozygous variant with a variety of mutation sites, leading to the occurrence of MLD disease.^34^ The homozygous mutation site c.925G>A was first found by Martina in a 1-year-old and a 6-month-old girl.^10^ The homozygous mutation site c.925G>A found in this study is the first such report in Asia.

It has been established that highly conserved amino acid sequences have functional value and are important for the protein structure, which suggests that they play a key role in determining the conformation of different domains of a protein. Missense mutation in genes can lead to changes in protein structure and spatial conformation by encoding amino acid substitutions. To observe the effect of c.925G>A and other c.925G mutation site (c.925G>T, c.925G>C) on protein structure and spatial conformation, we conducted a literature review of c.925G site mutations, compared ARSA and c.925G multi-species protein sequences and 3D structural model analysis.

The results were compiled according to a literature review of c.925G site mutations (c.925G>A, c.925G>T, c.925G>C) in which a compound heterozygous mutation or homozygous mutation causes MLD disease. The c.925G encodes the lysine at position 307 of the ARSA protein (p.E307), which is highly conserved among different species. It is indicated that amino acid changes at this position will significantly affect the structure and function of the ARSA protein. To explore the potential pathogenicity of the mutation sites found in this study, we used several tools to evaluate the pathogenicity of c.925G (c.925G>A, c.925G>T, c.925G>C). Multiple mutation evaluation procedures, SIFT, Polyphen-2, and MutationTaster, were used. The pathogenicity predictions consistently indicated that all three mutations at p.E307 are detrimental. The crystal structure of human arylsulfatase A shows that the core of the enzyme is composed of two β-pleated sheets. The major β-pleated sheet is formed by ten β-strands and is sandwiched between three α-helices on one side and four on the other.^41^ In our study, the 3D structural model analysis showed that p.E307 in the formation of a disulfide bond. Previous studies have described other pathogenic mutations (p.K304R, p.K304N, p.T306M, p.Y308H, p.E309Q, p.E309*, p.G310D, p.G311S, p.R313Q) in the disulfide bond region of ARSA,^10^ which may cause MLD. Our result suggests that p.E307 (p.E307K, p.E307Q, p.E307*) mutation may led to a change in the spatial steric hindrance and conformation of the ARSA structure, which may change the spatial location and morphology of the arylsulfatase A protein. Combined with literature review, multiple sequence alignment and protein structure analysis, it was found that the mutations of p.E307 (p.E307K, p.E307Q, p.E307*) may have pathogenic effects on ARSA protein structure.

To further elucidate the pathogenesis of MLD disease, we transferred a wild-type and mutant vector (c.925G>A, c.925G>T, c.925G>C) into the cell, and designed a MLD missense mutant cell model. This cell mutation model approach has been successfully applied to several rare disease studies.^42^ To more systematically analyze the potential effects of mutations on intracellular molecular processes, we obtained RNA-seq methods for transcriptome information from wild-type and mutant transfected cells. Based on transcriptome information, we further applied multi-dimensional system biology analysis using WGCNA combined with Hub gene screening and PPI analysis.

WGCNA is a systematic biological method for describing gene association patterns among different samples. It can be used to identify highly synergistic gene sets based on the coherence of gene sets and the association between gene sets and phenotypes. We applied WGCNA to construct a co-expression network for exploring mutations into three different bases at c.925G (c.925G>A, c.925G>T, c.925G>C). Functional enrichment analysis of ARSA mutation c.925G>A showed enrichment in pathways of calcium ion-dependent vesicle fusion, intra-Golgi vesicle-mediated transport, metaion-binding and ion transport. In the ARSA mutation c.925G>C, the enriched pathways were associated with the mitochondrial inner membrane, vesicle-mediated transport, and calcium binding. In the ARSA mutation c.925G>T, enrichment in the carbohydrate metabolism process pathway was found.

Hub genes with high connectivity may have more important biological significance in modules.^43^ The hub genes affected by the three mutations of c.925G provided functional support for the WGCNA results. The c.925G>A module identified hub genes related to Golgi vesicle-mediated transport (such as TRAPPC6B, COG5), cell junctions (SUN1), and meta ion-binding related genes (such as ZNF583, TRAPPC6B). The c.925G>C module identified hub genes related to the mitochondrial inner membrane (LRMP, SMIM24), vesicle-mediated transport (F5, LRMP, CALCR, AAK1, etc.), calcium binding (F5, CALCR). The c.925G>T module hub genes were enriched in the carbohydrate metabolism process pathway (BCL2L1, PSMC5).

PPI network analysis is an effective tool for revealing the principles of cellular organizations and functions. In this study, PPI analysis showed that the mutation of c.925G (c.925G>A, c.925G>T, c.925G>C) had more protein complexes that might interact than the control group. The composition of these protein complexes was mostly related to energy metabolism, ion binding and other signalling pathways.

The above functional analysis showed that the three mutations of c.925G had a comprehensive effect on cell function, including energy metabolism, ion binding, vesicle transport and ion transport (figure S3). Mutations in the ARSA gene result in the production of defective arylsulfatase A. As a result, inactive arylsulfatase A is transported to the lysosome, where it is unable to degrade various sulfate esters. This leads to sulfate-ester accumulation and subsequently to sulfatase deficiency. More than one catastrophic model was enriched with calcium-related functions, suggesting calcium channel-related drugs could be used to prevent progression of MLD disease. These results provide a potential pathogenic mechanism of ASRA mutation in metachromatic leukodystrophy and suggest possible strategies for diagnosis and treatment.

In conclusion, we identified a late infantile metachromatic leukodystrophy patient carrying on c.925G>A homozygous mutation in ARSA gene. According to literature and protein structure analysis, all mutation types of c.925G (c.925G>A, c.925G>T, c.925G>C) in ARSA gene are pathogenic. To analyze the contribution of mutations located on this site, we constructed overexpression cell models of wild type ARSA and c.925G mutation genes (c.925G>A, c.925G>T, c.925G>C). RNA-seq and bioinformatics results indicate that the mutations at c.925G cause comprehensive molecular changes related to energy metabolism, ion binding, vesicle transport and transport. These results provided potential pathogenic mechanism of ASRA mutation in metachromatic leukodystrophy and suggested possible ways in diagnosis and treatment.

## Acknowledgments

This work was supported by CAS Key Laboratory of Mental Health, Institute of Psychology.

The authors thank the participating individuals for their cooperation and their efforts in collecting the genetic information and DNA specimens.

