## Supplemental Table 1,2,3 4 and Supplementary figure 1,2,3,4 for "Identification of a missense ARSA mutation in metachromatic leukodystrophy and its potential pathogenic mechanism"

**Supplementary Table 1.** Quality of the Whole exome sequencing in this study.

| **Statistic** | **Sample** | | | |
| --- | --- | --- | --- | --- |
|  | I-1 | I-2 | II-1 | II-2 |
| Raw Bases (bp) | 17642855700 | 19467058800 | 21535325400 | 24746674200 |
| Clean Bases (bp) | 16399403923 | 18545867724 | 20188868376 | 23250333561 |
| Clean Q30 (%) | 92.48 | 94.9 | 94.93 | 94.64 |
| Clean Max GC/AT Bias Rate (%) | 7.69/6.88 | 7.30/7.44 | 7.71/6.60 | 7.60/6.91 |
| Clean GC Content (%) | 51 | 50 | 51 | 51 |
| Clean Reads Rate (%) | 92.95 | 95.27 | 93.75 | 93.95 |
| Capture Efficiency (%) | 62.25 | 57.18 | 63.78 | 63.1 |
| Clean Reads Mapped Rate (%) | 99.95 | 99.93 | 99.97 | 99.97 |
| Insert Size of Clean Reads (bp) | 256 | 267 | 242 | 244 |
| Insert Size Std Dev | 59.92 | 68.05 | 52.55 | 55.62 |
| Duplication Clean Reads Rate | 2.33 | 2.62 | 2.4 | 2.66 |
| Uniq Reads Rate (%) | 88.14 | 88.58 | 88.14 | 88.37 |
| Mean Depth of Target Region (X) | 102.74 | 103.72 | 124.75 | 138.26 |
| Coverage of Bases≥ 1x on Target Region (%) | 94 | 93.8 | 93.9 | 94 |
| Coverage of Bases≥ 20x on Target Region (%) | 92.5 | 92.6 | 92.8 | 93 |

Variations were using UCSC hg19 refGene (<http://genome.ucsc.edu/>);

The proband was the numbers II-2. The parents of proband was the number I-1 (father) and I-2 (mother). The healthy older brother of proband was the number II-1.

**Supplementary Table 2.** SNP Detection and Annotation in this study.

| **Statistics** | **Sample** | | | |
| --- | --- | --- | --- | --- |
|  | I-1 | I-2 | II-1 | II-2 |
| Total number of variants | 49674 | 49901 | 50420 | 51535 |
| Number of SNPs on target | 45396 | 45494 | 45887 | 46845 |
| Number of known dbSNP sites on target | 45145 | 45241 | 45629 | 46586 |
| Synonymous SNV | 9845 | 9828 | 9837 | 9975 |
| Non synonymous SNV | 8663 | 8590 | 8592 | 8785 |
| Stopgain | 68 | 63 | 60 | 71 |
| Stoploss | 8 | 7 | 10 | 7 |
| Exonic | 19994 | 19909 | 19947 | 20305 |
| Intronic | 21367 | 21553 | 21840 | 22343 |
| Splice site | 64 | 60 | 65 | 64 |
| 5'UTR | 983 | 972 | 1006 | 1020 |
| 3'UTR | 1256 | 1269 | 1282 | 1324 |
| ncRNA exonic | 969 | 939 | 962 | 1001 |
| ncRNA intronic | 944 | 966 | 972 | 1022 |
| ncRNA splicing | 2 | 2 | 2 | 1 |
| Upstream | 340 | 342 | 350 | 369 |
| Downstream | 160 | 165 | 173 | 163 |
| Intergenic | 1268 | 1254 | 1257 | 1291 |

Variations were using UCSC hg19 refGene (<http://genome.ucsc.edu/>).

The proband was the numbers II-2. The parents of proband was the number I-1 (father) and I-2 (mother). The healthy older brother of proband was the number II-1.

**Supplementary Table 3.** InDel Detection and Annotation in this study.

| **Statistics** | **Sample** | | | |
| --- | --- | --- | --- | --- |
|  | I-1 | I-2 | II-1 | II-2 |
| Total number of InDels | 4278 | 4407 | 4533 | 4690 |
| Ins-coding | 140 | 142 | 152 | 150 |
| Del-coding | 158 | 155 | 158 | 157 |
| 5'UTR | 113 | 106 | 122 | 117 |
| 3'UTR | 187 | 191 | 197 | 214 |
| Intergenic | 108 | 115 | 119 | 126 |
| Total insertion | 140 | 142 | 152 | 150 |
| Total deletion | 158 | 155 | 158 | 157 |
| Heterozygous InDels | 2656 | 2746 | 2832 | 2968 |
| Homozygous InDels | 1622 | 1661 | 1701 | 1722 |

Variations were using UCSC hg19 refGene (<http://genome.ucsc.edu/>);

The proband was the numbers II-2. The parents of proband was the number I-1 (father) and I-2 (mother). The healthy older brother of proband was the number II-1.

**Supplementary Table 4.** Characteristics of potential pathogenic mutations identified in this study.

| **Gene** | **Sample** | **Location** | **SNP ID** | **Nucleotide change** | **Predicted effect** | **depth/ Allele State** | **Type / region / protein effects** | **1000G/1000G.E/ESP/Exac/Exac.E/AD/ AD_EAS** | **SIFT/PP2/ MT** |
| --- | --- | --- | --- | --- | --- | --- | --- | --- | --- |
| ARSA | I-1 | chr22:51064636 | rs199476360 | c.925G>A | p.E309K | 207/ Heterozygosity | SNP/ exonic / nonsynonymous | NA/NA/NA/  NA/NA/0.000004139 /0.00005866 | D/D /D |
|  | I-2 | chr22:51064636 | rs199476360 | c.925G>A | p.E309K | 209/ Heterozygosity | SNP/ exonic / nonsynonymous | NA/NA/NA/  NA/NA/0.000004139 /0.00005866 | D/D /D |
|  | II-1 | - | - | - | - | - | - | - | - |
|  | II-2 | chr22:51064636 | rs199476360 | c.925G>A | p.E309K | 285/ Homozygosity | SNP/ exonic / nonsynonymous | NA/NA/NA/  NA/NA/0.000004139 /0.00005866 | D/D /D |
| SCN1A^*^ | I-1 | chr2:166911190 | rs777631884 | c.560G>A | p.R187Q | 31/ Heterozygosity | SNP/ exonic / nonsynonymous | NA/NA/NA  0.000008332//0.000004075 / | D/D /D |
|  | I-2 | - | - | - | - | - | - | - | - |
|  | II-1 | - | - | - | - | - | - | - | - |
|  | II-2 | chr2:166911190 | rs777631884 | c.560G>A | p.R187Q | 56 /Heterozygosity | SNP/ exonic / nonsynonymous | NA/NA/NA  0.000008332//0.000004075 / | D/D /D |
| SCN11^*^ | I-1 | - | - | - | - | - | - | - | - |
|  | I-2 | chr3:38888526 | rs143537709 | c.5035C>T | p.R1679 | 84/ Heterozygosity | SNP/ exonic / nonsynonymous | NA/NA/NA  0.00004123//0.00003257 /0.00005802 | D/D /D |
|  | II-1 | - | - | - | - | - | - | - | - |
|  | II-2 | chr3:38888526 | rs143537709 | c.5035C>T | p.R1679 | 114/ Heterozygosity | SNP/ exonic / nonsynonymous | NA/NA/NA  0.00004123//0.00003257 /0.00005802 | D/D /D |
| WNK1^*^ | I-1 | - | - | - | - | - | - | - | - |
|  | I-2 | chr12:1017956 | rs55650617 | c.7927T>A | p.X2643K | 70/ Heterozygosity | SNP\| exonic stoploss | 0.00179712/0.0089/NA  0.00008361/0.0012/0.00009761 /0.0014 | NA/NA /NA |
|  | II-1 | chr12:1017956 | rs55650617 | c.7927T>A | p.X2643K | 64/ Heterozygosity | SNP\| exonic stoploss | 0.00179712/0.0089/NA  0.00008361/0.0012/0.00009761 /0.0014 | NA/NA /NA |
|  | II-2 | chr12:1017956 | rs55650617 | c.7927T>A | p.X2643K | 67/ Heterozygosity | SNP\| exonic stoploss | 0.00179712/0.0089/NA  0.00008361/0.0012/0.00009761 /0.0014 | NA/NA /NA |

* Genetic variation detected did not match clinical manifestations；

Nucleotide numbers are derived from cDNA ARSA sequences, GenBank accession numbers: NM_000487.5; SCN1A GenBank accession number NM_001165963; SCN11 GenBank accession number NM_014139; WNK1GenBank accession number NM_001184985.

The symbol “–” indicates data not available.

SIFT, The SIFT score (dbNSFP version 3.0) indicates the effect of the mutation on the protein sequence with the value T or D. When the mutation affects multiple protein sequences at the same time, there is a SIFT value for each protein sequence, with the minimum value. D, deleterious (sift <=0.05); T, tolerated (sift > 0.05).

PolyPhen2, PolyPhen2 was used to predict the effect of the mutation on the protein sequence based on the HumanVar database for the diagnosis of Mendelian genetic diseases (dbNSFP version 3.0). The value is D or P or B. D, probably damaging (>=0.909); P, possibly damaging (0.447<=pp2_hvar<=0.909); B, benign (pp2_hvar<=0.446).

MutationTaster, MutationTaster predicted results (dbNSFP version 3.0) indicate the effect of the mutation on the protein sequence. Mutation Taster score, with a value of 0-1. The larger the score is the more reliable the prediction. MutationTaster predicted, with values of A, D, N or P. "A" ("Disease_causing_automatic"); D"("Disease_causing"); N" ("Polymorphism"); P"("Polymorphism_automatic"). Both A and D indicate that loci may be harmful.

NA, not applicable

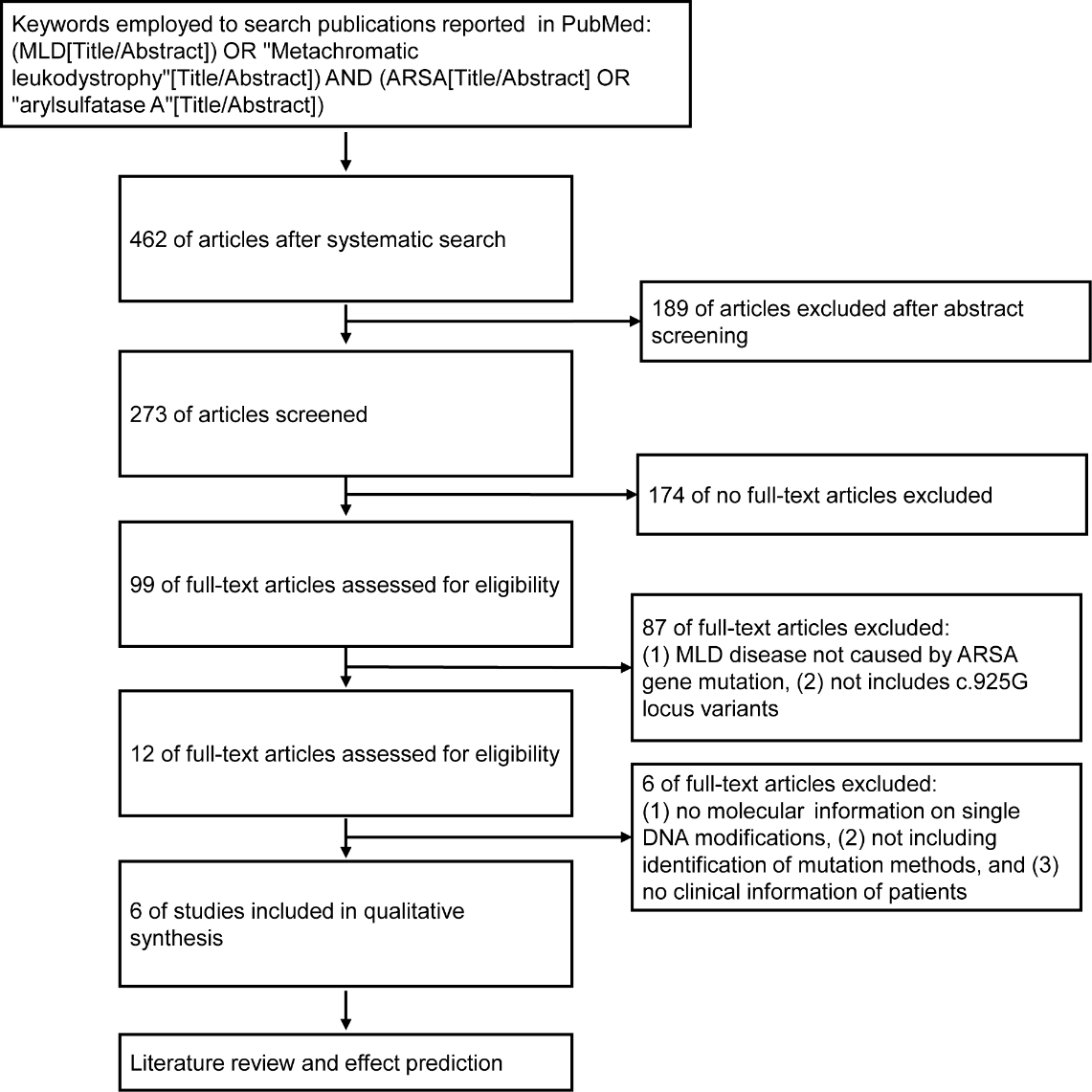

**Supplementary figure 1.** Flow of information through the different phases of a systematic literature review.

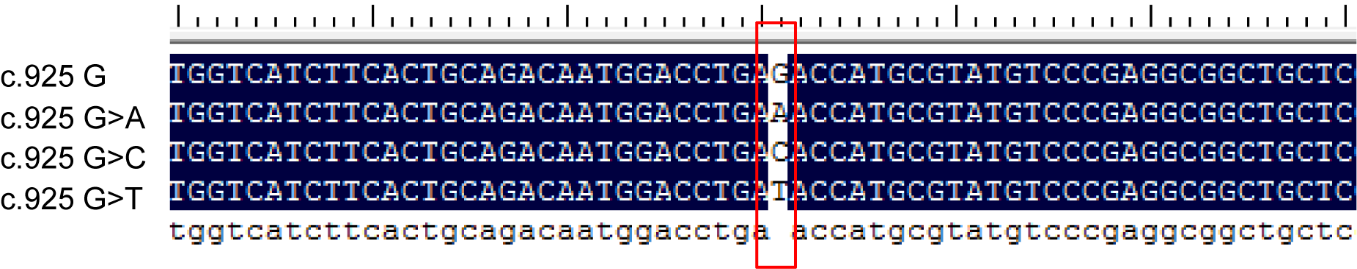

**Supplementary figure 2.** Wild-type ARSA cDNA /mutation vector sequences. The human wild-type ARSA cDNA cloned in the pCMV6 plasmid (c.925G). The three missense c.925G mutation (c.925G>A, c.925G>T, c.925G>C) were created on wild-type ARSA cDNA constructs, as shown in the red square. DNA sequences were confirmed using the ABI3500 sequencer. Nucleotide numbers are derived from cDNA ARSA sequences, GenBank accession numbers: NM_000487.5 and NP_000478.3.

**Supplementary Table 5.** Quality of the Transcriptomes in this study*

| **Sample** | **Raw reads** | **Raw Base (G)** | **Clean reads** | **Clean bases (G)** | **GC Content (%)** | **Q30**  **(%)** |
| --- | --- | --- | --- | --- | --- | --- |
| WT | 27075342 | 8.12 | 26156665 | 7.845 | 50.705 | 93.825 |
| vector | 25371499 | 7.61 | 24525447 | 7.355 | 50.925 | 93.63 |
| C.925 G | 25047944 | 7.515 | 24284098 | 7.285 | 52.13 | 93.925 |
| C.925 G>A | 26867676 | 8.06 | 26161371 | 7.85 | 52.14 | 93.845 |
| C.925 G>C | 23929807 | 7.18 | 23161745 | 6.95 | 52.115 | 93.935 |
| C.925 G>T | 26868186 | 8.06 | 26217328 | 7.865 | 52.095 | 93.47 |

*Three independent experiments with average values in the table.

**Supplementary Table 6.** Detection of hub genes and their functional annotations

| **Sample** | **Gene** | **GO Slim (Ontology)** | **Molecular Function (GO)** |
| --- | --- | --- | --- |
| c.925 G | HNF4G | [GO:0001071] nucleic acid binding transcription factor activity;[GO:0003674] molecular function;[GO:0005622] intracellular;[GO:0005575] cellular component;[GO:0003677] DNA binding;[GO:0004871] signal transducer activity;[GO:0043226] organelle;[GO:0005623] cell;[GO:0005634] nucleus;[GO:0005654] nucleoplasm;[GO:0043167] ion binding;[GO:0048856] anatomical structure development;[GO:0007165] signal transduction;[GO:0008150] biological process;[GO:0034641] cellular nitrogen compound metabolic process;[GO:0009058] biosynthetic process | GO:0003707 steroid hormone receptor activity;GO:0000978 RNA polymerase II proximal promoter sequence-specific DNA binding;GO:0000987 proximal promoter sequence-specific DNA binding |
|  | C10orf71 | NA | GO:0005515 protein binding;GO:0005488 binding;GO:0003674 molecular function |
|  | TIMM17A | [GO:0003674] molecular function;[GO:0005575] cellular component;[GO:0005622] intracellular;[GO:0055085] transmembrane transport;[GO:0008565] protein transporter activity;[GO:0022857] transmembrane transporter activity;[GO:0006605] protein targeting;[GO:0006810] transport;[GO:0008150] biological process;[GO:0007005] mitochondrion organization;[GO:0005737] cytoplasm;[GO:0005739] mitochondrion;[GO:0043234] protein complex;[GO:0005623] cell;[GO:0005634] nucleus;[GO:0005654] nucleoplasm;[GO:0043226] organelle | GO:0015450 P-P-bond-hydrolysis-driven protein transmembrane transporter activity;GO:0008320 protein transmembrane transporter activity;GO:1904680 peptide transmembrane transporter activity |
|  | IRF4 | [GO:0005829] cytosol;[GO:0006464] cellular protein modification process;[GO:0034641] cellular nitrogen compound metabolic process;[GO:0009058] biosynthetic process;[GO:0051276] chromosome organization;[GO:0008150] biological process;[GO:0002376] immune system process;[GO:0006950] response to stress;[GO:0007155] cell adhesion;[GO:0030154] cell differentiation;[GO:0048856] anatomical structure development;[GO:0007165] signal transduction;[GO:0000228] nuclear chromosome;[GO:0005575] cellular component;[GO:0005694] chromosome;[GO:0043226] organelle;[GO:0005622] intracellular;[GO:0005623] cell;[GO:0005634] nucleus;[GO:0001071] nucleic acid binding transcription factor activity;[GO:0003674] molecular function;[GO:0003677] DNA binding;[GO:0043234] protein complex;[GO:0008134] transcription factor binding;[GO:0008168] methyltransferase activity;[GO:0005737] cytoplasm;[GO:0005654] nucleoplasm | GO:0000978 RNA polymerase II proximal promoter sequence-specific DNA binding;GO:0000987 proximal promoter sequence-specific DNA binding;GO:0001228 DNA-binding transcription activator activity, RNA polymerase II-specific |
|  | SGCG | [GO:0005886] plasma membrane;[GO:0043226] organelle;[GO:0005856] cytoskeleton;[GO:0061024] membrane organization;[GO:0030154] cell differentiation;[GO:0043234] protein complex;[GO:0048856] anatomical structure development;[GO:0005575] cellular component;[GO:0005622] intracellular;[GO:0005737] cytoplasm;[GO:0005623] cell;[GO:0003674] molecular function;[GO:0008150] biological process;[GO:0003013] circulatory system process | GO:0005515 protein binding;GO:0005488 binding;GO:0003674 molecular function |
|  | EFCAB8 | [GO:0043167] ion binding;[GO:0003674] molecular function | GO:0005509 calcium ion binding;GO:0046872 metal ion binding;GO:0043169 cation binding |
|  | ST6GALNAC4 | [GO:0006464] cellular protein modification process;[GO:0005975] carbohydrate metabolic process;[GO:0005737] cytoplasm;[GO:0005794] Golgi apparatus;[GO:0043226] organelle;[GO:0005623] cell;[GO:0005575] cellular component;[GO:0005622] intracellular;[GO:0003674] molecular function;[GO:0016757] transferase activity, transferring glycosyl groups;[GO:0006629] lipid metabolic process;[GO:0008150] biological process;[GO:0034641] cellular nitrogen compound metabolic process;[GO:0009058] biosynthetic process | GO:0047290 (alpha-N-acetylneuraminyl-2,3-beta-galactosyl-1,3)-N-acetyl-galactosaminide 6-alpha-sialyltransferase activity;GO:0008373 sialyltransferase activity;GO:0016757 transferase activity, transferring glycosyl groups |
|  | SV2C | [GO:0003674] molecular function;[GO:0005623] cell;[GO:0005737] cytoplasm;[GO:0022857] transmembrane transporter activity;[GO:0055085] transmembrane transport;[GO:0016023] cytoplasmic membrane-bounded vesicle;[GO:0043226] organelle;[GO:0008150] biological process;[GO:0006810] transport;[GO:0005886] plasma membrane;[GO:0005575] cellular component;[GO:0005622] intracellular | GO:0022857 transmembrane transporter activity;GO:0005215 transporter activity;GO:0005515 protein binding |
|  | TMEM184A | [GO:0003674] molecular function;[GO:0006810] transport;[GO:0008150] biological process;[GO:0005622] intracellular;[GO:0005575] cellular component;[GO:0005623] cell;[GO:0005768] endosome;[GO:0043226] organelle;[GO:0005737] cytoplasm | GO:0008201 heparin binding;GO:0005539 glycosaminoglycan binding;GO:1901681 sulfur compound binding |
|  | PPP1R7 | [GO:0003674] molecular function;[GO:0005575] cellular component;[GO:0005622] intracellular;[GO:0005576] extracellular region;[GO:0008150] biological process;[GO:0030234] enzyme regulator activity;[GO:0006464] cellular protein modification process;[GO:0005634] nucleus;[GO:0043226] organelle;[GO:0005694] chromosome;[GO:0005737] cytoplasm;[GO:0005623] cell;[GO:0007059] chromosome segregation | GO:0019888 protein phosphatase regulator activity;GO:0019208 phosphatase regulator activity;GO:0030234 enzyme regulator activity |
|  | UTS2B | [GO:0005576] extracellular region;[GO:0005575] cellular component;[GO:0003674] molecular function;[GO:0003013] circulatory system process;[GO:0008150] biological process | GO:0005179 hormone activity;GO:0001664 G protein-coupled receptor binding;GO:0048018 receptor ligand activity |
|  | TRIM73 | [GO:0003674] molecular function;[GO:0005622] intracellular;[GO:0005575] cellular component;[GO:0043167] ion binding;[GO:0005623] cell | GO:0008270 zinc ion binding;GO:0046914 transition metal ion binding;GO:0046872 metal ion binding |
|  | SEPT12 | [GO:0043234] protein complex;[GO:0007049] cell cycle;[GO:0008150] biological process;[GO:0051301] cell division;[GO:0005575] cellular component;[GO:0005622] intracellular;[GO:0005623] cell;[GO:0005856] cytoskeleton;[GO:0043226] organelle;[GO:0003677] DNA binding;[GO:0003674] molecular function;[GO:0005737] cytoplasm;[GO:0043167] ion binding;[GO:0008289] lipid binding;[GO:0005929] cilium | GO:0019003 GDP binding;GO:0035091 phosphatidylinositol binding;GO:0003924 GTPase activity |
|  | CELA2A | [GO:0008233] peptidase activity;[GO:0003674] molecular function;[GO:0005623] cell;[GO:0005737] cytoplasm;[GO:0008150] biological process;[GO:0005576] extracellular region;[GO:0005615] extracellular space;[GO:0005622] intracellular;[GO:0005575] cellular component | GO:0004252 serine-type endopeptidase activity;GO:0008236 serine-type peptidase activity;GO:0017171 serine hydrolase activity |
| c.925 G>A | SUN1 | [GO:0003674] molecular function;[GO:0005575] cellular component;[GO:0005622] intracellular;[GO:0043226] organelle;[GO:0005635] nuclear envelope;[GO:0005634] nucleus;[GO:0005623] cell;[GO:0043234] protein complex;[GO:0007010] cytoskeleton organization;[GO:0008150] biological process;[GO:0061024] membrane organization;[GO:0005737] cytoplasm;[GO:0016023] cytoplasmic membrane-bounded vesicle;[GO:0007059] chromosome segregation;[GO:0051276] chromosome organization;[GO:0007049] cell cycle;[GO:0000003] reproduction | GO:0005521 lamin binding;GO:0043495 protein membrane anchor;GO:0005515 protein binding |
|  | ZNF583 | [GO:0003674] molecular function;[GO:0009058] biosynthetic process;[GO:0034641] cellular nitrogen compound metabolic process;[GO:0008150] biological process;[GO:0005622] intracellular;[GO:0005575] cellular component;[GO:0005623] cell;[GO:0003677] DNA binding;[GO:0001071] nucleic acid binding transcription factor activity;[GO:0005634] nucleus;[GO:0043226] organelle;[GO:0043167] ion binding | GO:0000981 DNA-binding transcription factor activity, RNA polymerase II-specific;GO:0003700 DNA-binding transcription factor activity;GO:0140110 transcription regulator activity |
|  | TRAPPC6B | [GO:0003674] molecular function;[GO:0005575] cellular component;[GO:0005622] intracellular;[GO:0005623] cell;[GO:0005737] cytoplasm;[GO:0005794] Golgi apparatus;[GO:0043226] organelle;[GO:0005975] carbohydrate metabolic process;[GO:0008150] biological process;[GO:0005829] cytosol;[GO:0005783] endoplasmic reticulum;[GO:0006464] cellular protein modification process;[GO:0022607] cellular component assembly;[GO:0065003] macromolecular complex assembly;[GO:0006461] protein complex assembly;[GO:0006810] transport;[GO:0016192] vesicle-mediated transport;[GO:0061024] membrane organization;[GO:0009058] biosynthetic process | GO:0005515 protein binding;GO:0005488 binding;GO:0003674 molecular function |
|  | SRSF6 | [GO:0003674] molecular function;[GO:0006397] mRNA processing;[GO:0008150] biological process;[GO:0022607] cellular component assembly;[GO:0022618] ribonucleoprotein complex assembly;[GO:0034641] cellular nitrogen compound metabolic process;[GO:0065003] macromolecular complex assembly;[GO:0003723] RNA binding;[GO:0009058] biosynthetic process;[GO:0006913] nucleocytoplasmic transport;[GO:0006810] transport;[GO:0005623] cell;[GO:0043226] organelle;[GO:0005654] nucleoplasm;[GO:0005634] nucleus;[GO:0008219] cell death;[GO:0006950] response to stress;[GO:0008283] cell proliferation;[GO:0048856] anatomical structure development;[GO:0030154] cell differentiation;[GO:0005622] intracellular;[GO:0005575] cellular component | GO:0036002 pre-mRNA binding;GO:0003723 RNA binding;GO:0003676 nucleic acid binding |
|  | CLTRN | [GO:0005575] cellular component;[GO:0005622] intracellular;[GO:0005576] extracellular region;[GO:0008150] biological process;[GO:0042592] homeostatic process;[GO:0006810] transport;[GO:0003674] molecular function;[GO:0005737] cytoplasm;[GO:0005623] cell;[GO:0061024] membrane organization;[GO:0016192] vesicle-mediated transport;[GO:0008233] peptidase activity;[GO:0043226] organelle;[GO:0007267] cell-cell signaling;[GO:0065003] macromolecular complex assembly;[GO:0022607] cellular component assembly;[GO:0006461] protein complex assembly;[GO:0005886] plasma membrane | GO:0042803 protein homodimerization activity;GO:0046983 protein dimerization activity;GO:0042802 identical protein binding |
|  | RIPK2 | [GO:0005622] intracellular;[GO:0005575] cellular component;[GO:0004871] signal transducer activity;[GO:0043167] ion binding;[GO:0005623] cell;[GO:0005737] cytoplasm;[GO:0043234] protein complex;[GO:0006914] autophagy;[GO:0008219] cell death;[GO:0009056] catabolic process;[GO:0008283] cell proliferation;[GO:0009058] biosynthetic process;[GO:0005829] cytosol;[GO:0005856] cytoskeleton;[GO:0043226] organelle;[GO:0034641] cellular nitrogen compound metabolic process;[GO:0007165] signal transduction;[GO:0008150] biological process;[GO:0003674] molecular function;[GO:0006464] cellular protein modification process;[GO:0016301] kinase activity;[GO:0007155] cell adhesion;[GO:0030154] cell differentiation;[GO:0048856] anatomical structure development;[GO:0002376] immune system process;[GO:0006950] response to stress | GO:0004706 JUN kinase kinase activity;GO:0089720 caspase binding;GO:0030274 LIM domain binding |
|  | DERL2 | [GO:0008150] biological process;[GO:0040007] growth;[GO:0003674] molecular function;[GO:0005575] cellular component;[GO:0005622] intracellular;[GO:0005737] cytoplasm;[GO:0005623] cell;[GO:0005768] endosome;[GO:0043226] organelle;[GO:0008283] cell proliferation;[GO:0006950] response to stress;[GO:0006810] transport;[GO:0007165] signal transduction;[GO:0009056] catabolic process;[GO:0009058] biosynthetic process;[GO:0006457] protein folding;[GO:0006464] cellular protein modification process;[GO:0005783] endoplasmic reticulum;[GO:0005975] carbohydrate metabolic process | GO:0005515 protein binding;GO:0005488 binding;GO:0003674 molecular function |
|  | ZBTB25 | [GO:0003674] molecular function;[GO:0043167] ion binding;[GO:0003677] DNA binding;[GO:0005575] cellular component;[GO:0005622] intracellular;[GO:0005623] cell;[GO:0005634] nucleus;[GO:0005737] cytoplasm;[GO:0043226] organelle;[GO:0005654] nucleoplasm;[GO:0001071] nucleic acid binding transcription factor activity;[GO:0009058] biosynthetic process;[GO:0008150] biological process;[GO:0034641] cellular nitrogen compound metabolic process | GO:0000981 DNA-binding transcription factor activity, RNA polymerase II-specific;GO:0003700 DNA-binding transcription factor activity;GO:0140110 transcription regulator activity |
|  | ZNF630 | [GO:0003674] molecular function;[GO:0008150] biological process;[GO:0034641] cellular nitrogen compound metabolic process;[GO:0009058] biosynthetic process;[GO:0005575] cellular component;[GO:0005623] cell;[GO:0005622] intracellular;[GO:0003677] DNA binding;[GO:0043226] organelle;[GO:0005634] nucleus;[GO:0043167] ion binding;[GO:0001071] nucleic acid binding transcription factor activity | GO:0003677 DNA binding;GO:0046872 metal ion binding;GO:0043169 cation binding |
|  | SLC30A8 | [GO:0005737] cytoplasm;[GO:0005623] cell;[GO:0005886] plasma membrane;[GO:0003674] molecular function;[GO:0043167] ion binding;[GO:0016023] cytoplasmic membrane-bounded vesicle;[GO:0043226] organelle;[GO:0007267] cell-cell signaling;[GO:0042592] homeostatic process;[GO:0016192] vesicle-mediated transport;[GO:0055085] transmembrane transport;[GO:0005575] cellular component;[GO:0022857] transmembrane transporter activity;[GO:0005622] intracellular;[GO:0006810] transport;[GO:0008150] biological process;[GO:0006950] response to stress;[GO:0002376] immune system process | GO:0005385 zinc ion transmembrane transporter activity;GO:0072509 divalent inorganic cation transmembrane transporter activity;GO:0046915 transition metal ion transmembrane transporter activity |
|  | CENPS-CORT | [GO:0003674] molecular function;[GO:0005576] extracellular region;[GO:0005575] cellular component | |
|  | ATP5MC2 | [GO:0005622] intracellular;[GO:0005575] cellular component;[GO:0005623] cell;[GO:0043226] organelle;[GO:0043234] protein complex;[GO:0005737] cytoplasm;[GO:0005739] mitochondrion;[GO:0008289] lipid binding;[GO:0022857] transmembrane transporter activity;[GO:0055085] transmembrane transport;[GO:0006810] transport;[GO:0006091] generation of precursor metabolites and energy;[GO:0008150] biological process;[GO:0034641] cellular nitrogen compound metabolic process;[GO:0044281] small molecule metabolic process;[GO:0009058] biosynthetic process;[GO:0003674] molecular function | GO:0046933 proton-transporting ATP synthase activity, rotational mechanism;GO:0044769 ATPase activity, coupled to transmembrane movement of ions, rotational mechanism;GO:0019829 cation-transporting ATPase activity |
|  | PHOSPHO2 | [GO:0003674] molecular function;[GO:0043167] ion binding;[GO:0008150] biological process;[GO:0016791] phosphatase activity | GO:0033883 pyridoxal phosphatase activity;GO:0016791 phosphatase activity;GO:0042578 phosphoric ester hydrolase activity |
|  | IL12B | [GO:0005623] cell;[GO:0005737] cytoplasm;[GO:0009058] biosynthetic process;[GO:0050877] neurological system process;[GO:0043234] protein complex;[GO:0044403] symbiosis, encompassing mutualism through parasitism;[GO:0040011] locomotion;[GO:0048870] cell motility;[GO:0040007] growth;[GO:0007165] signal transduction;[GO:0006810] transport;[GO:0006913] nucleocytoplasmic transport;[GO:0006605] protein targeting;[GO:0008219] cell death;[GO:0007049] cell cycle;[GO:0003674] molecular function;[GO:0002376] immune system process;[GO:0008283] cell proliferation;[GO:0006464] cellular protein modification process;[GO:0006950] response to stress;[GO:0007155] cell adhesion;[GO:0030154] cell differentiation;[GO:0048856] anatomical structure development;[GO:0000003] reproduction;[GO:0008150] biological process;[GO:0005575] cellular component;[GO:0005622] intracellular;[GO:0004871] signal transducer activity;[GO:0005576] extracellular region;[GO:0005615] extracellular space | GO:0042164 interleukin-12 alpha subunit binding;GO:0045519 interleukin-23 receptor binding;GO:0019972 interleukin-12 binding |
|  | COG5 | [GO:0005634] nucleus;[GO:0005654] nucleoplasm;[GO:0016192] vesicle-mediated transport;[GO:0008150] biological process;[GO:0006810] transport;[GO:0043234] protein complex;[GO:0003674] molecular function;[GO:0005829] cytosol;[GO:0005622] intracellular;[GO:0043226] organelle;[GO:0005623] cell;[GO:0005737] cytoplasm;[GO:0005794] Golgi apparatus;[GO:0005575] cellular component | GO:0005515 protein binding;GO:0005488 binding;GO:0003674 molecular function |
|  | MTCP1 | [GO:0005622] intracellular;[GO:0005575] cellular component;[GO:0005623] cell;[GO:0005737] cytoplasm;[GO:0043226] organelle;[GO:0005739] mitochondrion;[GO:0008283] cell proliferation;[GO:0008150] biological process | GO:0043539 protein serine/threonine kinase activator activity;GO:0030295 protein kinase activator activity;GO:0019209 kinase activator activity |
| c.925 G>C | CFAP126 | [GO:0003674] molecular function;[GO:0005623] cell;[GO:0005737] cytoplasm;[GO:0005886] plasma membrane;[GO:0005929] cilium;[GO:0005815] microtubule organizing center;[GO:0005856] cytoskeleton;[GO:0043226] organelle;[GO:0008150] biological process;[GO:0005622] intracellular;[GO:0005575] cellular component | |
|  | F5 | [GO:0043167] ion binding;[GO:0005615] extracellular space;[GO:0005576] extracellular region;[GO:0005575] cellular component;[GO:0005622] intracellular;[GO:0005623] cell;[GO:0005737] cytoplasm;[GO:0005794] Golgi apparatus;[GO:0043226] organelle;[GO:0008150] biological process;[GO:0006810] transport;[GO:0016192] vesicle-mediated transport;[GO:0003013] circulatory system process;[GO:0003674] molecular function;[GO:0009058] biosynthetic process;[GO:0061024] membrane organization;[GO:0006950] response to stress;[GO:0016023] cytoplasmic membrane-bounded vesicle;[GO:0005783] endoplasmic reticulum;[GO:0006464] cellular protein modification process;[GO:0022607] cellular component assembly;[GO:0006461] protein complex assembly;[GO:0065003] macromolecular complex assembly;[GO:0005886] plasma membrane;[GO:0005975] carbohydrate metabolic process | GO:0005507 copper ion binding;GO:0046914 transition metal ion binding;GO:0046872 metal ion binding |
|  | LRMP | [GO:0002376] immune system process;[GO:0005783] endoplasmic reticulum;[GO:0005815] microtubule organizing center;[GO:0005886] plasma membrane;[GO:0005575] cellular component;[GO:0005622] intracellular;[GO:0005623] cell;[GO:0005856] cytoskeleton;[GO:0043226] organelle;[GO:0008150] biological process;[GO:0000003] reproduction;[GO:0016192] vesicle-mediated transport;[GO:0061024] membrane organization;[GO:0006810] transport;[GO:0005634] nucleus;[GO:0005694] chromosome;[GO:0005635] nuclear envelope;[GO:0005737] cytoplasm | |
|  | MTRNR2L4 | [GO:0005737] cytoplasm;[GO:0005623] cell;[GO:0005575] cellular component;[GO:0005622] intracellular;[GO:0005576] extracellular region | GO:0048019 receptor antagonist activity;GO:0030547 receptor inhibitor activity;GO:0030545 receptor regulator activity |
|  | ELP4 | [GO:0005737] cytoplasm;[GO:0005622] intracellular;[GO:0005623] cell;[GO:0005634] nucleus;[GO:0005654] nucleoplasm;[GO:0043226] organelle;[GO:0043234] protein complex;[GO:0005575] cellular component;[GO:0003674] molecular function;[GO:0019899] enzyme binding;[GO:0016746] transferase activity, transferring acyl groups;[GO:0006464] cellular protein modification process;[GO:0008150] biological process;[GO:0030234] enzyme regulator activity;[GO:0051276] chromosome organization;[GO:0034641] cellular nitrogen compound metabolic process;[GO:0009058] biosynthetic process | GO:0008607 phosphorylase kinase regulator activity;GO:0000993 RNA polymerase II complex binding;GO:0043175 RNA polymerase core enzyme binding |
|  | IL5RA | [GO:0030154] cell differentiation;[GO:0048856] anatomical structure development;[GO:0000902] cell morphogenesis;[GO:0008150] biological process;[GO:0006464] cellular protein modification process;[GO:0007165] signal transduction;[GO:0005622] intracellular;[GO:0005615] extracellular space;[GO:0005576] extracellular region;[GO:0005575] cellular component;[GO:0004871] signal transducer activity;[GO:0005886] plasma membrane;[GO:0005623] cell;[GO:0002376] immune system process;[GO:0006950] response to stress;[GO:0003674] molecular function;[GO:0040011] locomotion;[GO:0008283] cell proliferation | GO:0004914 interleukin-5 receptor activity;GO:0004896 cytokine receptor activity;GO:0019955 cytokine binding |
|  | CALCR | [GO:0003674] molecular function;[GO:0004871] signal transducer activity;[GO:0008150] biological process;[GO:0007165] signal transduction;[GO:0005575] cellular component;[GO:0005622] intracellular;[GO:0005623] cell;[GO:0005737] cytoplasm;[GO:0008565] protein transporter activity;[GO:0016192] vesicle-mediated transport;[GO:0042592] homeostatic process;[GO:0006810] transport;[GO:0007009] plasma membrane organization;[GO:0061024] membrane organization;[GO:0044281] small molecule metabolic process;[GO:0034641] cellular nitrogen compound metabolic process;[GO:0009058] biosynthetic process;[GO:0005829] cytosol;[GO:0005886] plasma membrane;[GO:0016023] cytoplasmic membrane-bounded vesicle;[GO:0043226] organelle;[GO:0002376] immune system process;[GO:0048856] anatomical structure development;[GO:0030154] cell differentiation;[GO:0005929] cilium | GO:0032841 calcitonin binding;GO:0001635 calcitonin gene-related peptide receptor activity;GO:0097643 amylin receptor activity |
|  | AAK1 | [GO:0008150] biological process;[GO:0043167] ion binding;[GO:0016301] kinase activity;[GO:0006464] cellular protein modification process;[GO:0003674] molecular function;[GO:0016192] vesicle-mediated transport;[GO:0007165] signal transduction;[GO:0006810] transport;[GO:0043226] organelle;[GO:0016023] cytoplasmic membrane-bounded vesicle;[GO:0005622] intracellular;[GO:0005575] cellular component;[GO:0005886] plasma membrane;[GO:0005737] cytoplasm;[GO:0005623] cell | GO:0035612 AP-2 adaptor complex binding;GO:0005112 Notch binding;GO:0004674 protein serine/threonine kinase activity |
|  | SRSF9 | [GO:0005575] cellular component;[GO:0005622] intracellular;[GO:0005634] nucleus;[GO:0005623] cell;[GO:0043226] organelle;[GO:0005730] nucleolus;[GO:0005654] nucleoplasm;[GO:0003674] molecular function;[GO:0006397] mRNA processing;[GO:0008150] biological process;[GO:0022607] cellular component assembly;[GO:0022618] ribonucleoprotein complex assembly;[GO:0034641] cellular nitrogen compound metabolic process;[GO:0065003] macromolecular complex assembly;[GO:0003723] RNA binding;[GO:0006810] transport;[GO:0009058] biosynthetic process;[GO:0006913] nucleocytoplasmic transport | GO:0019904 protein domain specific binding;GO:0003723 RNA binding;GO:0003676 nucleic acid binding |
|  | SMIM24 | [GO:0043226] organelle;[GO:0008150] biological process;[GO:0003674] molecular function;[GO:0005575] cellular component;[GO:0005576] extracellular region | GO:0003674 molecular function |
|  | ZC3H12D | [GO:0004518] nuclease activity;[GO:0003674] molecular function;[GO:0040007] growth;[GO:0007049] cell cycle;[GO:0008150] biological process;[GO:0005575] cellular component;[GO:0005622] intracellular;[GO:0005623] cell;[GO:0005737] cytoplasm;[GO:0043226] organelle;[GO:0005634] nucleus;[GO:0043167] ion binding;[GO:0034641] cellular nitrogen compound metabolic process | GO:0004519 endonuclease activity;GO:0004518 nuclease activity;GO:0016788 hydrolase activity, acting on ester bonds |
|  | STT3A | [GO:0005575] cellular component;[GO:0009058] biosynthetic process;[GO:0006464] cellular protein modification process;[GO:0005975] carbohydrate metabolic process;[GO:0008150] biological process;[GO:0003674] molecular function;[GO:0016757] transferase activity, transferring glycosyl groups;[GO:0005622] intracellular;[GO:0005623] cell;[GO:0005737] cytoplasm;[GO:0005783] endoplasmic reticulum;[GO:0043226] organelle;[GO:0043234] protein complex | GO:0004579 dolichyl-diphosphooligosaccharide-protein glycotransferase activity;GO:0004576 oligosaccharyl transferase activity;GO:0016758 transferase activity, transferring hexosyl groups |
|  | ACTL6A | [GO:0005886] plasma membrane;[GO:0006464] cellular protein modification process;[GO:0051276] chromosome organization;[GO:0006259] DNA metabolic process;[GO:0006950] response to stress;[GO:0034641] cellular nitrogen compound metabolic process;[GO:0009058] biosynthetic process;[GO:0007165] signal transduction;[GO:0040007] growth;[GO:0006810] transport;[GO:0006605] protein targeting;[GO:0048856] anatomical structure development;[GO:0005575] cellular component;[GO:0005622] intracellular;[GO:0005623] cell;[GO:0005634] nucleus;[GO:0005654] nucleoplasm;[GO:0043226] organelle;[GO:0043234] protein complex;[GO:0008150] biological process;[GO:0009056] catabolic process;[GO:0006914] autophagy;[GO:0007005] mitochondrion organization;[GO:0005694] chromosome;[GO:0000228] nuclear chromosome;[GO:0000988] transcription factor activity, protein binding;[GO:0003674] molecular function;[GO:0003677] DNA binding | GO:0000980 RNA polymerase II distal enhancer sequence-specific DNA binding;GO:0031492 nucleosomal DNA binding;GO:0001158 enhancer sequence-specific DNA binding |
|  | IFIH1 | [GO:0005575] cellular component;[GO:0005622] intracellular;[GO:0043167] ion binding;[GO:0003674] molecular function;[GO:0008219] cell death;[GO:0044403] symbiosis, encompassing mutualism through parasitism;[GO:0005829] cytosol;[GO:0006464] cellular protein modification process;[GO:0005737] cytoplasm;[GO:0005623] cell;[GO:0005634] nucleus;[GO:0043226] organelle;[GO:0008150] biological process;[GO:0002376] immune system process;[GO:0006950] response to stress;[GO:0007165] signal transduction;[GO:0003723] RNA binding;[GO:0003677] DNA binding;[GO:0004386] helicase activity | GO:0003725 double-stranded RNA binding;GO:0003727 single-stranded RNA binding;GO:0004386 helicase activity |
| c.925 G>T | BCL2L1 | [GO:0006810] transport;[GO:0061024] membrane organization;[GO:0007009] plasma membrane organization;[GO:0009056] catabolic process;[GO:0006950] response to stress;[GO:0016192] vesicle-mediated transport;[GO:0006914] autophagy;[GO:0016023] cytoplasmic membrane-bounded vesicle;[GO:0044403] symbiosis, encompassing mutualism through parasitism;[GO:0019899] enzyme binding;[GO:0043234] protein complex;[GO:0005575] cellular component;[GO:0005622] intracellular;[GO:0007049] cell cycle;[GO:0008150] biological process;[GO:0051301] cell division;[GO:0005623] cell;[GO:0005737] cytoplasm;[GO:0043226] organelle;[GO:0005635] nuclear envelope;[GO:0005739] mitochondrion;[GO:0005634] nucleus;[GO:0002376] immune system process;[GO:0003674] molecular function;[GO:0007005] mitochondrion organization;[GO:0007165] signal transduction;[GO:0008219] cell death;[GO:0005829] cytosol;[GO:0005815] microtubule organizing center;[GO:0005856] cytoskeleton;[GO:0008283] cell proliferation;[GO:0030154] cell differentiation;[GO:0040007] growth;[GO:0000003] reproduction;[GO:0009790] embryo development;[GO:0048856] anatomical structure development | GO:0051434 BH3 domain binding;GO:0051400 BH domain binding;GO:0070513 death domain binding |
|  | S100A4 | [GO:0005575] cellular component;[GO:0043167] ion binding;[GO:0005576] extracellular region;[GO:0005615] extracellular space;[GO:0005622] intracellular;[GO:0008150] biological process;[GO:0048856] anatomical structure development;[GO:0030154] cell differentiation;[GO:0000902] cell morphogenesis;[GO:0003723] RNA binding;[GO:0003674] molecular function;[GO:0043226] organelle;[GO:0005634] nucleus;[GO:0005623] cell;[GO:0005737] cytoplasm;[GO:0007165] signal transduction;[GO:0008092] cytoskeletal protein binding | GO:0050786 RAGE receptor binding;GO:0048306 calcium-dependent protein binding;GO:0003779 actin binding |
|  | PSMC5 | [GO:0003674] molecular function;[GO:0043167] ion binding;[GO:0008134] transcription factor binding;[GO:0009056] catabolic process;[GO:0006520] cellular amino acid metabolic process;[GO:0016887] ATPase activity;[GO:0044403] symbiosis, encompassing mutualism through parasitism;[GO:0008219] cell death;[GO:0040011] locomotion;[GO:0002376] immune system process;[GO:0006464] cellular protein modification process;[GO:0007165] signal transduction;[GO:0005575] cellular component;[GO:0005622] intracellular;[GO:0005623] cell;[GO:0043234] protein complex;[GO:0000902] cell morphogenesis;[GO:0048856] anatomical structure development;[GO:0030154] cell differentiation;[GO:0000988] transcription factor activity, protein binding;[GO:0007049] cell cycle;[GO:0008150] biological process;[GO:0006950] response to stress;[GO:0044281] small molecule metabolic process;[GO:0034641] cellular nitrogen compound metabolic process;[GO:0022607] cellular component assembly;[GO:0006461] protein complex assembly;[GO:0065003] macromolecular complex assembly;[GO:0009058] biosynthetic process;[GO:0005829] cytosol;[GO:0005576] extracellular region;[GO:0005615] extracellular space;[GO:0005737] cytoplasm;[GO:0005634] nucleus;[GO:0043226] organelle;[GO:0005654] nucleoplasm | GO:0031531 thyrotropin-releasing hormone receptor binding;GO:0051428 peptide hormone receptor binding;GO:0017025 TBP-class protein binding |
|  | GNG11 | [GO:0006091] generation of precursor metabolites and energy;[GO:0008150] biological process;[GO:0005886] plasma membrane;[GO:0043234] protein complex;[GO:0005623] cell;[GO:0004871] signal transducer activity;[GO:0005575] cellular component;[GO:0005622] intracellular;[GO:0007165] signal transduction;[GO:0044281] small molecule metabolic process;[GO:0003674] molecular function;[GO:0003924] GTPase activity | GO:0031681 G-protein beta-subunit binding;GO:0003924 GTPase activity;GO:0017111 nucleoside-triphosphatase activity |
|  | OTUD5 | [GO:0008233] peptidase activity;[GO:0008150] biological process;[GO:0006464] cellular protein modification process;[GO:0003674] molecular function;[GO:0006950] response to stress;[GO:0005829] cytosol;[GO:0005737] cytoplasm;[GO:0005623] cell;[GO:0005622] intracellular;[GO:0005575] cellular component;[GO:0002376] immune system process | GO:0061578 Lys63-specific deubiquitinase activity;GO:1990380 Lys48-specific deubiquitinase activity;GO:0004843 thiol-dependent ubiquitin-specific protease activity |
|  | CLEC16A | [GO:0005764] lysosome;[GO:0005768] endosome;[GO:0006914] autophagy;[GO:0008150] biological process;[GO:0043226] organelle;[GO:0005575] cellular component;[GO:0005622] intracellular;[GO:0005623] cell;[GO:0005737] cytoplasm;[GO:0005773] vacuole;[GO:0003674] molecular function | GO:0017137 Rab GTPase binding;GO:0017016 Ras GTPase binding;GO:0031267 small GTPase binding |
|  | CCDC116 | [GO:0003674] molecular function;[GO:0005622] intracellular;[GO:0005575] cellular component;[GO:0005815] microtubule organizing center;[GO:0005856] cytoskeleton;[GO:0043226] organelle;[GO:0005737] cytoplasm;[GO:0005623] cell | GO:0005515 protein binding;GO:0005488 binding;GO:0003674 molecular function |
|  | NOP53 | [GO:0008150] biological process;[GO:0003674] molecular function;[GO:0003723] RNA binding;[GO:0005634] nucleus;[GO:0043226] organelle;[GO:0005730] nucleolus;[GO:0005623] cell;[GO:0005575] cellular component;[GO:0005622] intracellular | GO:0008097 5S rRNA binding;GO:0002039 p53 binding;GO:0019843 rRNA binding |

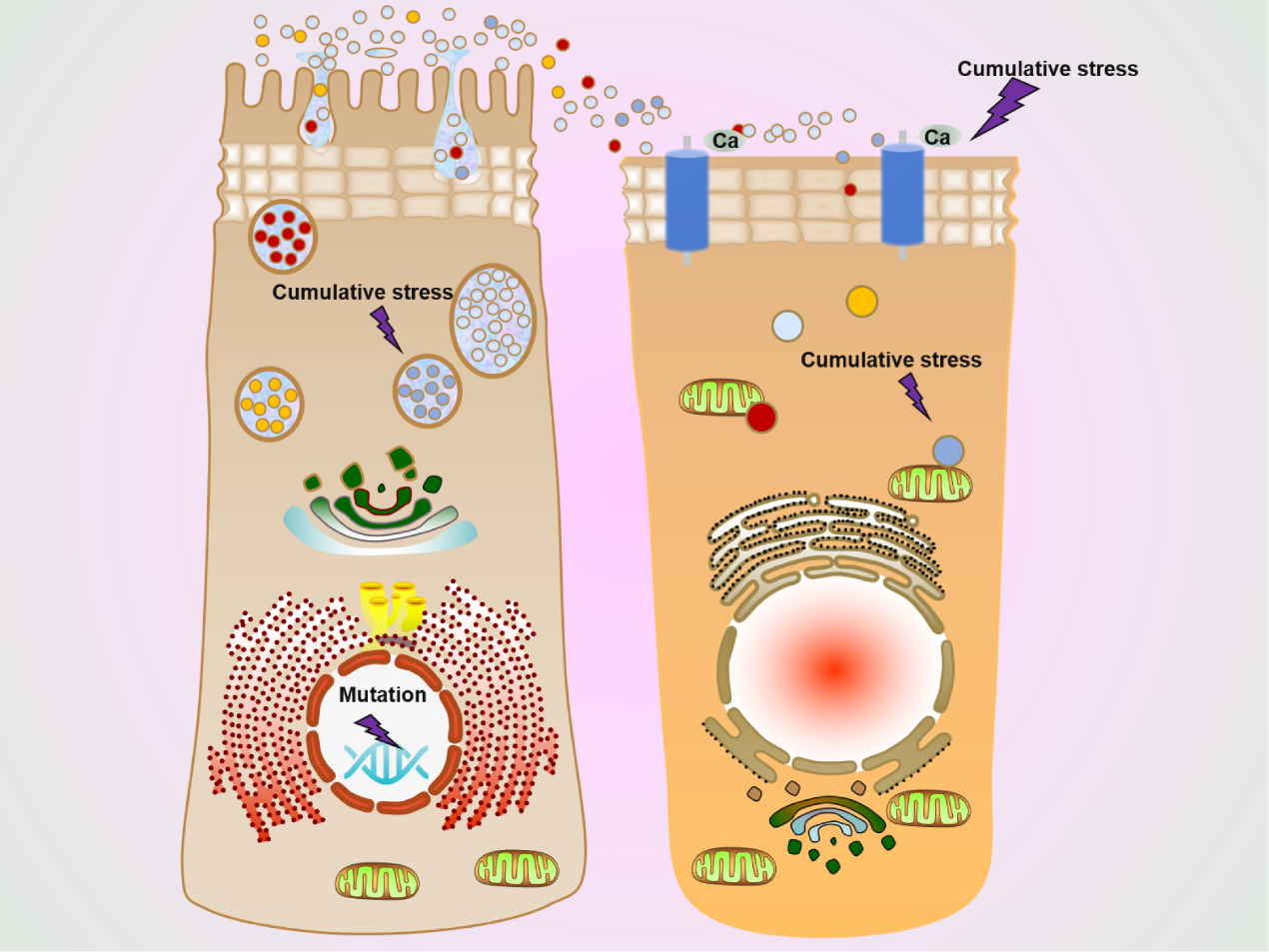

**Supplementary figure 3.** The defect in arylsulfatase A deficiency. ARSA gene c.925G three mutation (c. 925G>A, c. 925G>T, c. 925G>C) result in the production of defective the arylsulfatase A enzyme. It is speculated that the mutation in this region will cause the change of disulfide bond, lead to accumulation pressure, and affect the calcium ion binding and the structure of mitochondria and other related organelle membranes.
